# Structural basis of LRPPRC-SLIRP-dependent translation by the mitoribosome

**DOI:** 10.1101/2022.06.20.496763

**Authors:** Vivek Singh, J. Conor Moran, Yuzuru Itoh, Iliana C. Soto, Flavia Fontanesi, Mary Couvillion, Martijn A. Huynen, Stirling Churchman, Antoni Barrientos, Alexey Amunts

**Author notes:** These authors contributed equally (Vivek Singh, J. Conor Moran).

## Abstract

In mammalian mitochondria, mRNAs are co-transcriptionally stabilized by the protein factor LRPPRC. Here, we characterize LRPPRC as an mRNA delivery factor and report its cryo-EM structure in complex with SLIRP, mRNA and the mitoribosome. The structure shows that LRPPRC associates with the mitoribosomal proteins mS39 and the N-terminus of mS31 through recognition of the LRPPRC helical repeats. Together, the proteins form a corridor for hand-off the mRNA. The mRNA is directly bound to SLIRP, which also has a stabilizing function for LRPPRC. To delineate the effect of LRPPRC on individual mitochondrial transcripts, we used an RNAseq approach, metabolic labeling and mitoribosome profiling that showed a major influence on ND1, ND2, ATP6, COX1, COX2, and COX3 mRNA translation efficiency. Our data suggest that LRPPRC-SLIRP acts in recruitment of mitochondrial mRNAs to modulate their translation. Collectively, the data define LRPPRC-SLIRP as a regulator of the mitochondrial gene expression system.

---

The mitoribosome is organised in a small and large subunit (SSU and LSU) that are assembled from multiple components in a coordinated manner and through regulated sequential mechanisms ^1–5^. The SSU formation is accomplished by the association of the mitoribosomal protein mS37 and the initiation factor mtIF3, leading to a mature state that is ready for translation of the mRNA ^3, 6^. In mammals, mitochondrial transcription is polycistronic and gives rise to two long transcripts, corresponding to almost the entire heavy and light mtDNA strands. The individual mRNAs are available for translation only after they are liberated from the original polycistronic transcripts and polyadenylated ^7^. In *Escherichia coli*, a functional transcription-translation coupling mechanism has been characterised involving a physical association of the RNA polymerase with the SSU, termed the expressome ^8–10^. In contrast, in mammalian mitochondria, nucleoids are not compartmented with protein synthesis; mitoribosomes are independently tethered to the membrane ^11, 12^, and no coupling with the RNA polymerase has been reported. In addition, human mitochondrial mRNAs and the mitoribosome do not have the Shine–Dalgarno (SD) and anti-SD sequences that are used to recruit mRNA to SSU in bacteria ^13^. Mitochondrial mRNAs also lack cap 5′ modifications, which is a hallmark of eukaryotic cytosolic mRNAs translation initiation. In the cytosol, mRNA is recruited to a pre-initiation complex, consisting of the SSU and translation initiation factors, which then scans along the 5′ untranslated region to find the start codon ^14, 15^. No equivalent mechanism has been found in mitochondria, and thus, how mRNAs are delivered for translation in mitochondria remained unknown.

The 130-kDa protein factor LRPPRC (leucine-rich pentatricopeptide repeat-containing protein), a member of a Metazoa-specific pentatricopeptide repeat family, was reported to act as a global mitochondrial mRNA chaperone that binds co-transcriptionally ^16–18^. LRPPRC is an integral part of the post-transcriptional processing machinery required for mRNA stability, polyadenylation, and translation ^16–19^. Mutations in the gene encoding for LRPPRC lead to French-Canadian type Leigh syndrome (LSFC) an untreatable paediatric neurodegenerative disorder caused by ultimately impaired mitochondrial energy conversion ^20^.

LRPPRC has been reported to interact with a small 11-kDa protein cofactor SLIRP (SRA stem-loop-interacting RNA-binding protein) ^21, 22^ that plays roles in LRPPRC stability and maintaining steady-state mRNA levels ^23^. *SLIRP* silencing results in the destabilization of respiratory complexes, loss of enzymatic activity, and reduction in mRNA levels, implicating a role in mRNA homeostasis ^24^. *SLIRP* variants cause a respiratory deficiency that leads to mitochondrial encephalomyopathy ^25^. In addition, *SLIRP* knockdown results in increased turnover of LRPPRC ^23, 25, 26^, and *in vivo* co-stabilisation suggests that the two entities have interdependent functions ^23, 27^. The interaction of LRPPRC and SLIRP *in vitro* has been previously studied ^28^.

LRPPRC has also been implicated in coordinating mitochondrial mRNA stability and translation ^18, 29^. Previous analysis showed a correlation between presence of LRPPRC and mRNA on the mitoribosome ^30^. However, there are no structures available for LRPPRC, SLIRP, or any complexes containing them, and *in vitro* reconstitution could not provide meaningful information, in part because not all the components of the mitochondrial gene expression system have been characterised. Thus, although isolated mitoribosomal models have been determined ^31–,33^, the molecular mechanisms of mRNA delivery to the SSU for activation of translation and the potential involvement of LRPPRC-SLIRP in this process remained unknown.

## Results

### Structure determination of LRPPRC-SLIRP bound to the mitoribosome

To explore the molecular basis for translation activation in human mitochondria, we used low salt conditions to isolate a mitoribosome:LRPPRC-SLIRP-mRNA complex for cryo-EM. We merged particles containing tRNAs in the A- or P-site, as well as an extra density in the vicinity of the mRNA entry channel and applied iterative local-masked refinement and classification with signal subtraction (Extended Data Fig. 1a). It resulted in a 2.9 Å resolution map of the mitoribosome during mRNA delivery to the SSU, with the local resolution for the LRPPRC binding region of ∼3.4 Å (Extended Data Fig. 1b,c). The reconstruction showed a clear density only for the LRPPRC N-terminal domains (residues 64-644, average local resolution ∼4.5 Å) bound to the SSU head, which is consistent with a previous mass-spectrometry analysis (Extended Data Fig. 1d,e) ^30^. It allowed us to model 34 α-helices, 17 of which (α2-18) form a ring-like architecture, while the rest form an extended tail that adopts a 90° curvature and projects 110 Å from the SSU body in parallel to the L7/L12 stalk (Fig. 1a,b). The C-terminal domains (residues 645-1394) were not resolved. The complete LRPPRC model obtained with *AlphaFold2* ^34^ combined with Translation/Liberation/Screw Motion Determination (TLSMD) analysis ^35, 36^ defined the C-terminal domains as individual segments, indicating potential flexibility (Extended Data Fig. 2).

**Figure 1.**
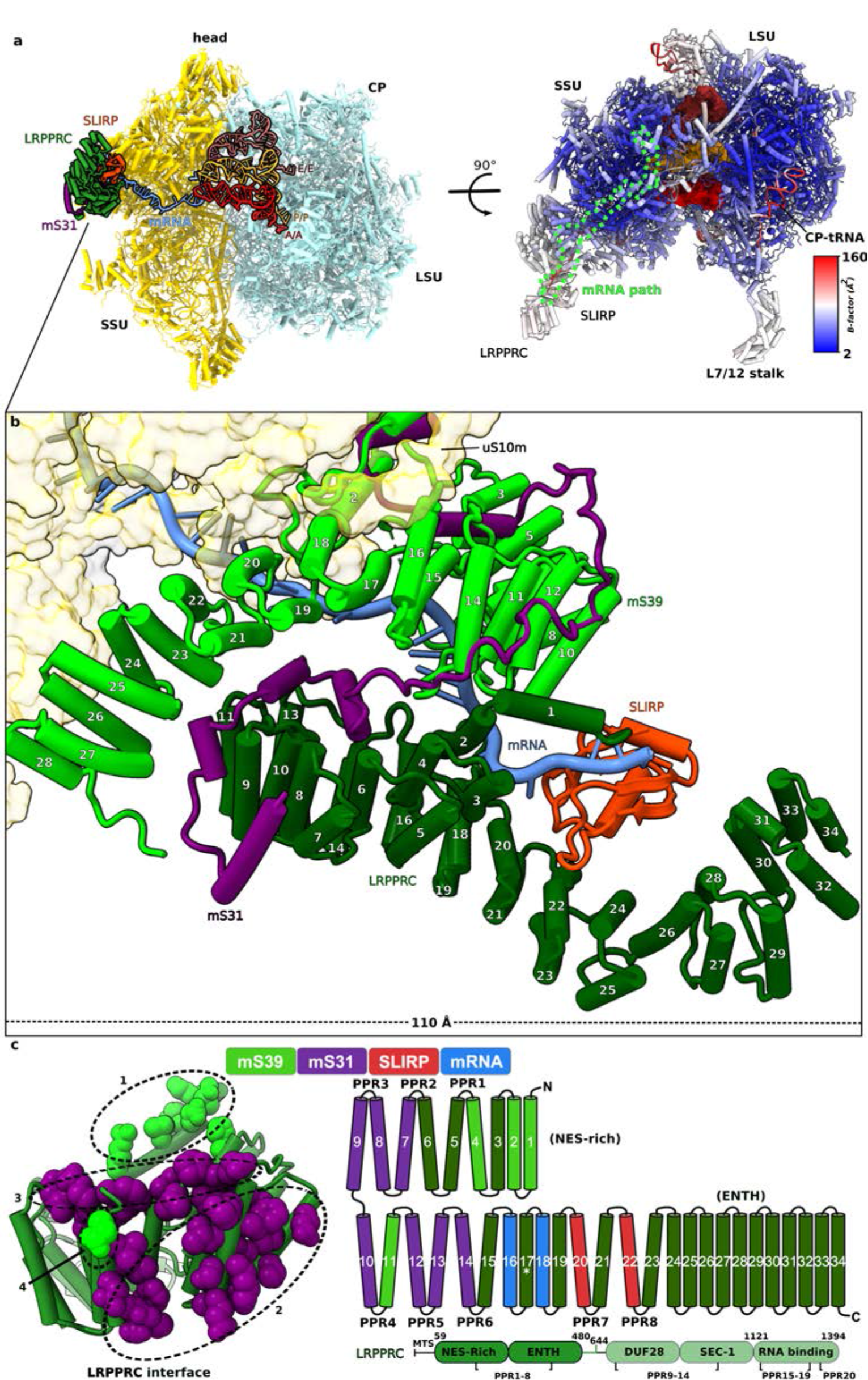
Structure of mitoribosome with LRPPRC-SLIRP bound to mRNA. **a,** Overview of the mitoribosome:LRPPRC-SLIRP model. Right panel, top view of the model colored by atomic *B*-factor (Å^2^), tRNAs in surface (red, orange, brown), mRNA path (light green) is highlighted. **b,** A close-up view of the mitoribosome:LRPPRC-SLIRP-mRNA interactions. LRPPRC associates with mS31-m39 via a ring-like structure (α2-18) that together form a corridor for the hand-off the mRNA from SLIRP. **c,** Contact sites between LRPPRC and mS31-mS39 (within 4 Å distance), view from the interface. Right panel, schematic diagram showing the topology of LRPPRC consisting of 34 helices. Colours represent engagement in interactions with mS39 (light green), mS31 (purple), SLIRP (orange), mRNA (blue). The position of LSFC variant (A354V) is indicated with an asterisk on helix 17.

**Figure 2.**
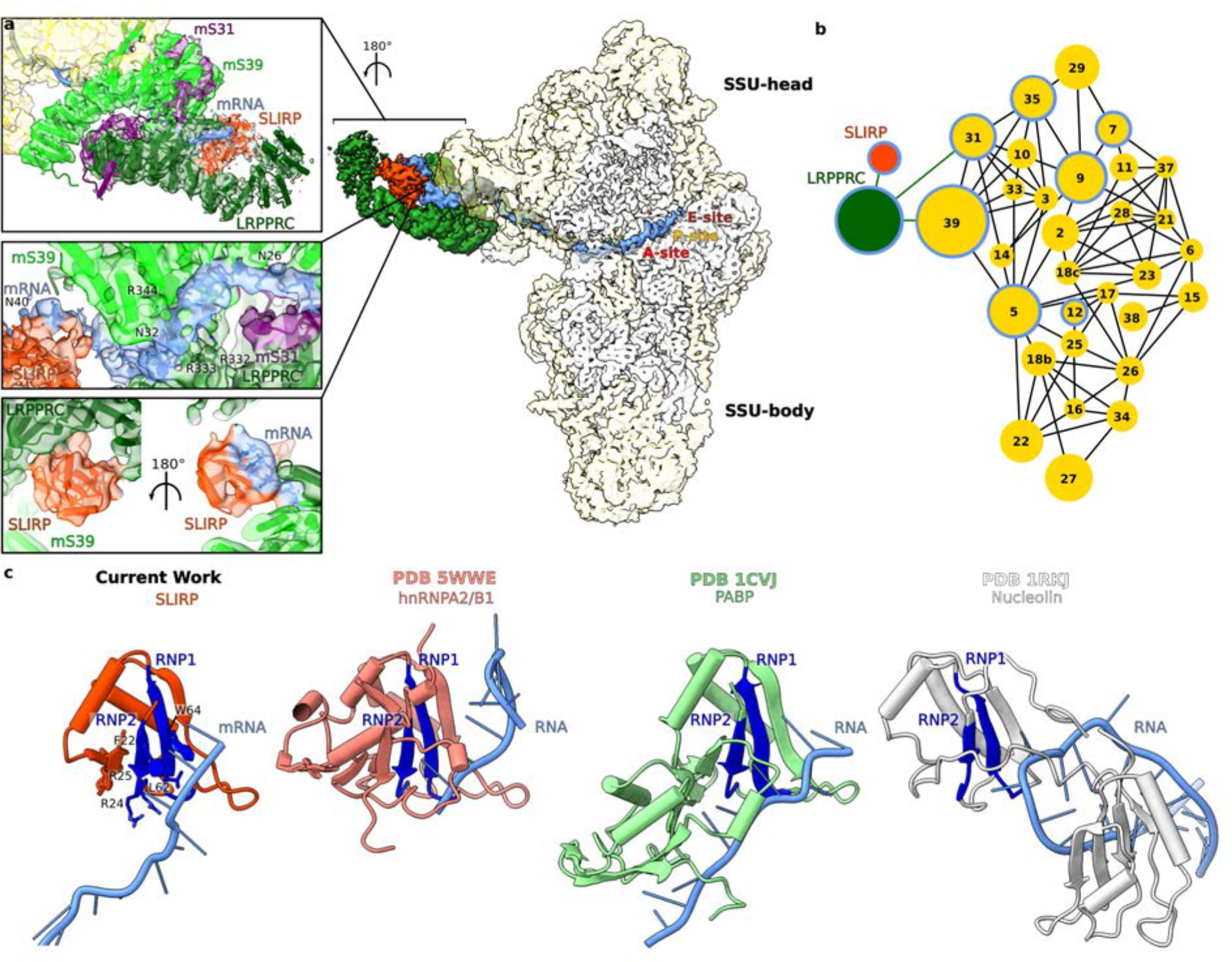
Overview of density for LRPPRC, SLIRP and mRNA and their interactions with SSU proteins. **a,** The density map for LRPPRC (dark green), SLIRP (orange), mRNA (blue) on the SSU is shown in the centre. The model and map for mS39-LRPPRC-SLIRP and corresponding bound mRNA residues are shown in closeup views on the left, and arginines involved in mRNA binding are indicated. The bottom closeup views show SLIRP with its associated densities for LRPPRC and mRNA. For clarity, the map for SLIRP has been low-pass filtered to 6 Å resolution. **b,** Schematic of protein-protein interactions, where node size corresponds to relative molecular mass. Nodes of proteins involved in mRNA binding are encircled in blue. **c,** RRM containing proteins: SLIRP, hnRNPA1/B2 (PDBID 5WWE), PolyA binding protein (PABP, PDBID 1CVJ) and Nucleolin (PDBID 1RKJ) are shown in complex with RNA with RNP1 and RNP2 sub-motifs colored blue.

When LRPPRC-SLIRP is bound to the mitoribosome, a previously disordered density of mS31 that extends from the core also becomes ordered, revealing its N-terminal region (Fig. 2a). This region is arranged in two helix-turn-helix motifs, offering a 1930 Å^2^ surface area for direct interactions with LRPPRC (Fig. 1c, Fig. 2). The position of the LRPPRC residue 354, in which the mutation A354V leads to LSFC with a clinically distinct cytochrome *c* oxidase deficiency and acute fatal acidotic crises is in a buried area of helix 17, close to the mRNA binding region (Fig. 1c, Extended Data Fig. 2a). A previous study demonstrated that the mutation is abolishing the interaction with the protein SLIRP ^37^. Consistent with mass spectrometry analysis ^28^ and the interaction interface previously determined ^37^, the remaining associated density was assigned as SLIRP, found to be located close to the Epsin N-terminal homology (ENTH) domain of LRPPRC (Fig. 2a). Finally, SLIRP is connected to an elongated density on the LRPPRC surface that is also associated with six of the mitoribosomal proteins and corresponds to the endogenous mRNA (Fig. 2).

### SLIRP is stably associated with mRNA and LRPPRC on the SSU

The binding of SLIRP in our model is enabled via LRPPRC helices α20 and 22, which is consistent with crosslinking mass spectrometry data and mutational analysis ^28^. The structure reveals that SLIRP links the nuclear export signal (NES) domain with the curved region of the ENTH domain of LRPPRC (Fig. 1b, Fig. 1c). This binding of SLIRP contributes to a corridor for the mRNA that extends to mS31 and mS39 (Fig. 1b, Supplementary Video 1). Through this corridor, the mRNA extends over ∼180 Å all the way to the decoding center (Fig. 2a). In our structure, SLIRP is oriented such that the conserved RNA recognition motif (RRM), including its submotifs RNP1 (residues 21-26) and RNP2 (residues 60-67) ^38–40^ form an interface with modeled mRNA (Fig. 2). The arrangement of RNP1 and RNP2 with respect to the mRNA is similar to that observed in previously reported structures of other RRM proteins ^41–43^ (Fig. 2c, Extended Data Fig. 3). Moreover, residues R24, R25 of RNP2 and L62 of RNP1 motifs previously implicated to be required for RNA binding by SLIRP ^37^ are positioned within an interacting distance of the mRNA (Fig. 2a). Thus, SLIRP contributes to the LRPPRC specific scaffold, and accounts for a role in binding the mRNA.

**Figure 3.**
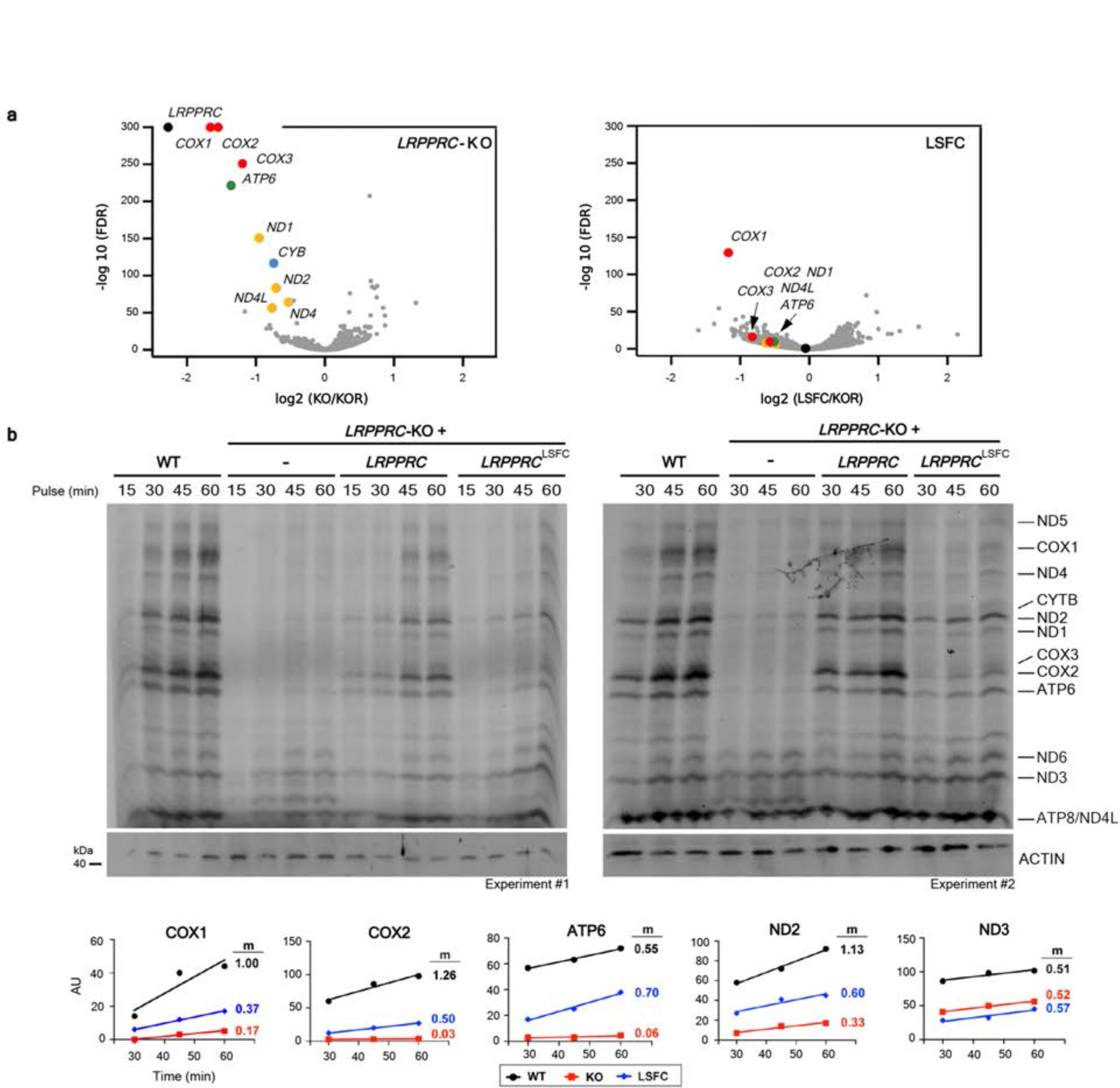
Mechanism of LRPPRC-SLIRP mediated mRNA binding and stabilization. **a,** Whole-cell RNAseq normalized by read depth, comparing *LRPPRC*-KO cells (KO) with KO cells reconstituted with WT *LRPPRC* (KOR) or the LSFC variant (A354V). The results are the average of two biological replicates. The differentially expressed mitochondrial transcripts are color-coded: coding for subunits of cytochrome *c* oxidase (CIV) in red, NADH dehydrogenase (CI) in yellow, coenzyme Q-cytochrome *c* oxidoreductase (CIII) in blue, and ATP synthase (CV) in green. **b,** Metabolic labeling with [^35^S]-labeled methionine of newly-synthesized mitochondrial polypeptides for the indicated times, in the presence of emetine to inhibit cytosolic protein synthesis, in whole HEK-293T WT, *LRPPRC-*KO cells, and KO cells reconstituted with LRPPRC (KO+WT) or the LSFC variant (KO+LSFC). Bottom panel, representative plots of [^35^S]-labeled methionine incorporation into specific polypeptides in WT or *LRPPRC*-KO cells.

The *B*-factor distribution of SLIRP in our model is similar to that of LRPPRC, while still lower than some of the more mobile components of the mitoribosome, such as the acceptor arm of the CP-tRNA^Val^ (Fig. 1a). This indicates a functionally relevant association with LRPPRC in terms of stability of binding. Our finding that SLIRP is involved in hand-off the mRNA to the mitoribosome provides a mechanistic explanation for the previous results from biochemical studies showing that SLIRP affects LRPPRC properties *in vitro* ^27, 28^, and the presentation of the mRNA to the mitoribosome *in vivo* ^23^.

Since in *E. coli*, the expressome-mediating protein NusG was proposed to regulate mRNA unwinding ^8^, and SSU proteins uS3 and uS4 have an intrinsic RNA helicase activity ^44^, we searched for known helicase signature motifs ^45^ in the LRPPRC sequence, but no such motifs were present. In the mitoribosome, where the mRNA channel entry site is located, a bacteria-like ring-shaped entrance is missing, the entrance itself has shifted, and its diameter expanded ^31^. The mRNA extends all the way into the head/beak of the SSU stabilised by mitoribosome-specific components: mS39 helical repeats, mS35 N-terminus that extends from the side of the SSU head, and N-terminal extension of uS9m that contacts the mRNA nucleotide at position 15 (numbered from the E-site).

### LRPPRC-SLIRP hands-off the mRNA to mS31-mS39, channeling it for translation

Next, we analysed the structural basis for the complex formation. The association of LRPPRC with the mitoribosome involves the helices α1, 2, 6-11, that form a mitoribosome-binding surface (Fig. 1b, Fig. 1c). The binding is mediated by four distinct contacting regions (Extended Data Fig. 6): 1) α1-2 (residues 64-95) is flanked by a region of mS39, a PPR domain-containing protein, that consists of four bundled helices (α11-14); 2) α7 and 9 form a shared bundle with two N-terminal helices of mS31 (residues 175-208, stabilized by C-terminal region of mS39) that encircle the NES-rich domain to complement the PPR domain; 3) α10-11 are capped by a pronounced turn of mS31 (residues 209-232) acting as a lid that marks the LRPPRC boundary, and it is sandwiched by the mS39 helix α19 and C-terminus from the opposite side; 4) in addition, α11 is also positioned directly against the helix α23 of mS39. Thus, LRPPRC docks onto the surface of the mitoribosome via mS31-mS39, which are tightly associated with each other, and each provides two contact patches to contribute to stable binding.

Based on the structural analysis, the hand-off of the mRNA for translation is mediated by four of the LRPPRC helices: α1, 2, 16, and 18 (Fig. 1b, Fig. 1c, Supplementary Video 1). The mRNA nucleotides 33-35 (numbered from the E-site) are stabilized in a cleft formed by α1-2 on one side and α16, 18 on the other. Nucleotides 31 and 32 contact residues R332, R333 from LRPPRC, as well as R344 from mS39 (Fig. 2a). This region is within 120-130 Å from the P-site. The involvement of the NES-rich domain of LRPPRC in mRNA binding in our structure is consistent with a biochemical analysis of recombinant LRPPRC where the N-terminal PPR segments were systematically removed, which showed a reduced formation of protein-RNA complexes ^28^. The rest of the mRNA is situated too far from LRPPRC to interact with it. Here, the mRNA is handed to mS31-mS39, consistently with a translation initiation complex ^46^.

In the structure, mS31, mS39, and LRPPRC together form a 60-Å long corridor that channels the mRNA from SLIRP toward the mitoribosomal core (Fig. 1b, Supplementary Video 1). The binding of LRPPRC is mediated by four distinct contact regions (Extended Data Fig. 6): contact- 1 is formed by α1-2 (residues 64-95) flanked by the mS39 PPR domain of four bundled helices (α11-14); contact-2 is formed by α7, 9 that generate a shared bundle with two N-terminal helices of mS31 (residues 175-208, stabilized by C-terminal region of mS39) that encircle the head to complement the PPR domain; contact-3 is formed by α10-11 that are capped by a pronounced turn of mS31 (residues 209-232) acting as a lid that marks the LRPPRC boundary, and it is sandwiched by the mS39 helix α21 and C-terminus from the opposite side; contact-4 is formed by α11 that is positioned directly against the mS39 α23.

With respect to mRNA binding, nucleotides 26-30 bind mS39 PPR domain 5, and nucleotide 26 connects to contact-2 (Fig. 2a, Extended Data Fig. 6). Thus, the mRNA hand-off is achieved through functional cooperation between LRPPRC-SLIRP and mS31-mS39. Therefore, in the mitoribosome:LRPPRC-SLIRP with mRNA model, LRPPRC performs three functions: coordination of SLIRP, which plays a key role in the process of mRNA recruitment, association with the SSU, and hand-off of the mRNA for translation (Fig. 1, Extended Data Fig. 6). Supplementary Video 1).

### LRPPRC is recruited for translation of mRNAs

Next, we asked whether LRPRRC-SLIRP delivers all mRNAs to the mitoribosome or is selective. We generated an *LRPPRC*-knockout cell line ^29^ that was rescued with either a wild-type *LRPPRC* or a variant carrying the LSFC founder mutation A354V ^20^ (Extended Data Fig. 4). The steady-state levels of the LSFC variant were reduced by 60%, suggesting protein instability as reported in patients ^22^, and the levels of SLIRP were equally decreased (Extended Data Fig. 4). We then implemented an RNAseq approach that confirmed a substantially depleted mitochondrial transcriptome ^17, 47, 48^ (Fig. 3a). In the LRPPRC-knockout, transcripts from the heavy strand were lowered by 1.5-4-fold, except for ND3, which remained stable, consistent with protein synthesis data (Fig. 3b), while the single light strand-encoded *ND6* mRNA was not affected as reported ^16^, and the effect of the LSFC mutation on RNA stability was limited to six transcripts (Fig. 3a), indicating LRPPRC’s role in heavy strand mRNA stability is non-specific. Metabolic labeling assays using [^35^S]- methionine indicated that incorporation of the radiolabeled amino acid into most newly synthesized mitochondrial proteins is severely decreased in *LRPPRC*-KO cells (Fig. 3b). However, there were differential effects among transcripts; synthesis of ND3, ND4L, and ATP8 remained above 50% of the WT, but translation of other transcripts proceeded at a lower rate (*e.g*., *ND1*, *ND2*, or *ATP6*) or was virtually blocked (*e.g*., *COX1, COX2, or COX3)* (Fig. 3b). The translational defect results in a decrease in the steady-state levels of the four OXPHOS complexes that contain mtDNA-encoded subunits (Extended Data Fig. 5).

**Figure 4.**
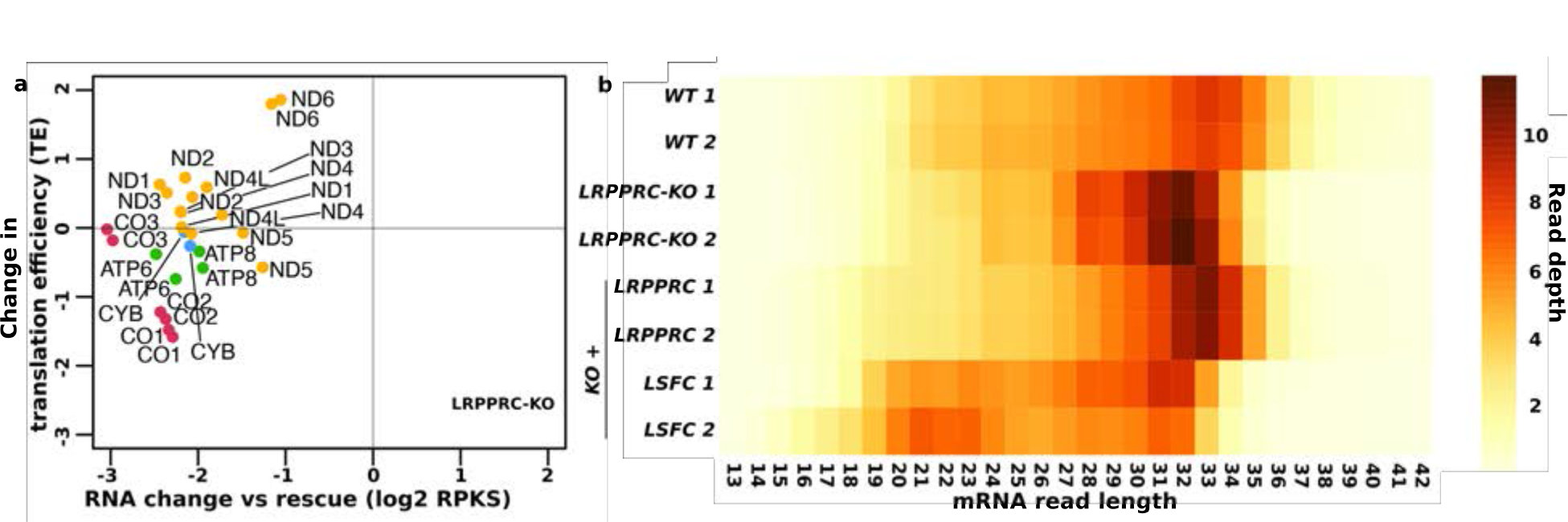
**Mitochondrial translation efficiency is decreased in *LRPPRC*-KO cells. a**, Change in TE and RNA abundance in *LRPPRC*-KO cells compared to LRPPRC-reconstituted cells (“rescue”). Mitoribosome profiling data and RNA-seq data were normalized using a mouse lysate spike-in control (RPKS) ^29^, then TE was calculated from these normalized values (mitoribosome profiling / RNA-seq). Biological replicates are shown as individual points. The mitochondrial transcripts are color-coded as in Fig. 3. **b**, Heat map showing the length distribution for reads mapping to mitochondrial mRNAs ^29^.

**Figure 5.**
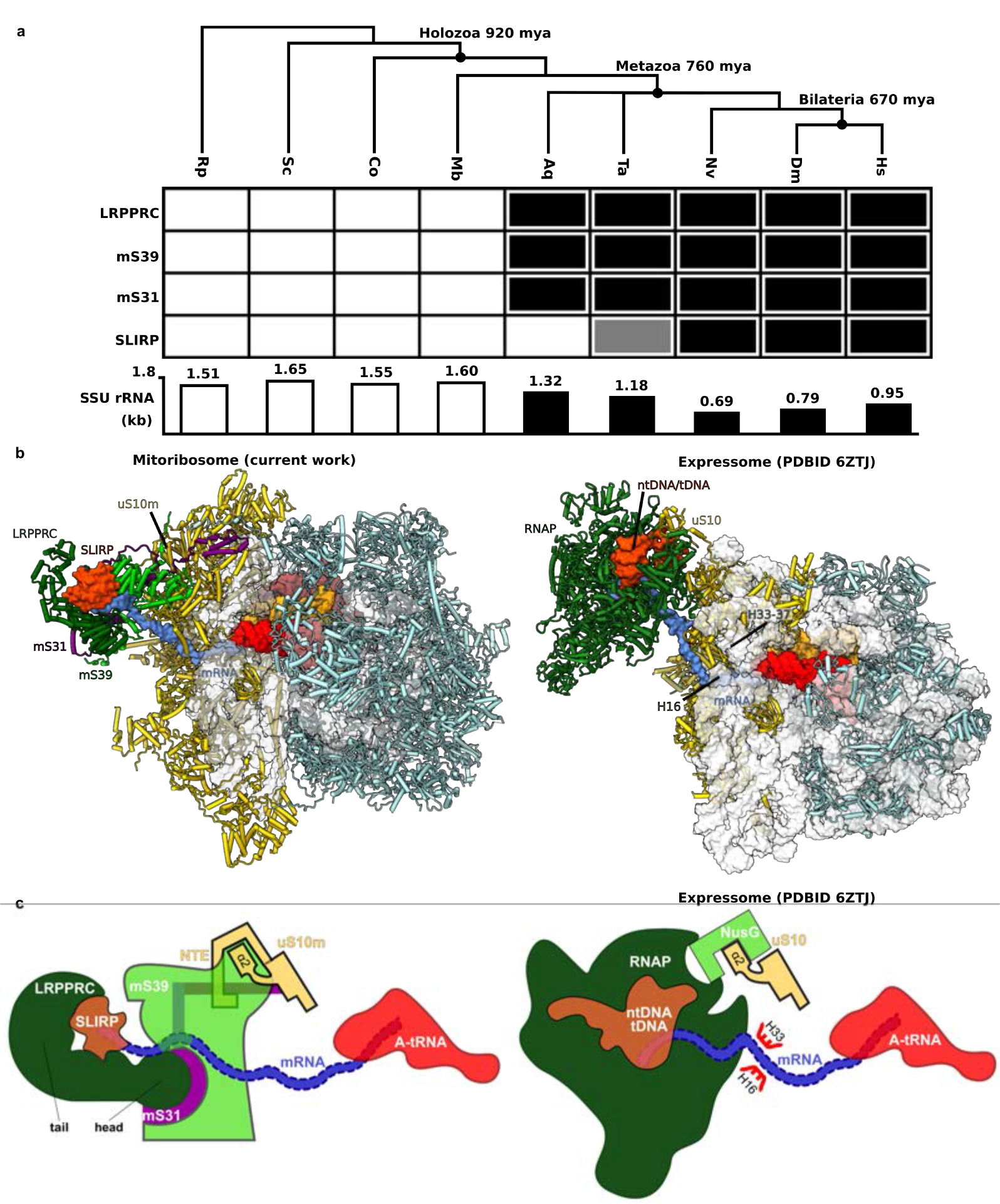
Formation of mitoribosome:LRPPRC-SLIRP and 70S:RNAP complexes. **a,** Phylogenetic analysis shows correlation between acquisition of LRPPRC, SLIRP, mS31, mS39 and reduction of rRNA in Metazoa. Black rectangles indicate the presence of proteins, grey indicates uncertainty about the presence of an ortholog. *Hs*: *Homo sapiens, Ds: Drosophila melanogaster, Nv*: *Nematostella vectensis, Ta: Trichoplax adhaerens, Aq*: *Amphimedon queenslandica* (sponge)*, Mb*: *Monosiga brevicollis* (unicellular choanoflagellate)*, Co*: *Capsaspora owczarzaki* (protist)*, Sc*: *Saccharomyces cerevisiae* (fungi)*, Rp*: *Rickettsia prowazekii* (alpha-proteobacterium). Dating in million years ago (mya) is based on previously described method ^55^. **b,** Model of the mitoribosome:LRPPRC-SLIRP complex compared with the uncoupled model of the expressome from *E. coli* ^8^. **c,** Schematic representation indicating association of mRNA-delivering proteins in the mitoribosome compared to NusG-coupled expressome ^8^.

To assess whether the mitochondrial translation efficiency (TE) is decreased in an mRNA- specific manner in the *LRPPRC*-knockout, we performed mitoribosome profiling (Fig. 4). These data show a decreased TE, calculated by dividing spike-in normalized ribosome footprint reads by the spike-in normalized RNA sequencing reads (in other words, how well a particular transcript is translated) ^49, 50^. In *LRPPRC-*knockout cells, TE was attenuated for *COX1* and *COX2* transcripts and the bicistronic *ATP8/ATP6* transcript, intriguingly more for ATP6 than ATP8 (Fig. 4a). Thus, our data suggest that LRPPRC-SLIRP is required for the translation of some transcripts, in agreement with the metabolic labeling of newly synthesized mitochondrial products (Fig. 3b).

To support the role of LRPPRC in mRNA binding, we determined an average length of the mitoribosome-protected fragments using mitoribosome profiling (Fig 4b). In the *LRPPRC*- knockout cells, we observed a decrease in the average protected fragment length, compared to the wild type (Fig 4b). This observation is consistent with the structural data showing the association of LRPPRC with mRNA and the mitoribosome. Previous studies also showed that LRPPRC–SLIRP relaxes structures of mRNAs ^17^, potentially to expose it to initiate translation ^16^. The average protected fragment length in LSFC cells were smaller than in wild-type cells, similar to the *LRPPRC*-knockout (Fig 4b), suggesting that whereas the mutant protein participates in translation, it does so differently than the wild-type protein

The images were quantified in two independent experiments.

### The mitoribosome:LRPPRC-SLIRP complex is specific to Metazoa

To place the structural data into an evolutionary context, we performed comparative phylogenetic analysis of the proteins involved in the mRNA hand-off process. Since the mitochondrial rRNA has been generally reduced in Metazoa ^51^, we examined whether this loss might coincide with the origin of LRPPRC and its interactors. The orthology database EGGNOGG ^52^ and previous analysis ^53^ indicated that LRPPRC and mS31 are only present in Bilateria, while mS39 only occurs in Metazoa. We then confirmed the results with more sensitive homology detection ^54^ followed by manual sequence analysis examining domain composition, which put the origin of LRPPRC and mS31 at the root of the Metazoa. Thus, the appearance of these proteins coincides with the loss of parts of the rRNA (Fig. 5a). SLIRP appears to originate slightly later than LRPPRC, but its small size makes determining its phylogenetic origin less conclusive.

The correlation between rRNA reduction and protein acquisition is important, because the rRNA regions h16 (410-432) and h33-37 (997-1118) that bridge the mRNA to the channel entrance in bacteria ^8^ are either absent or reduced in the metazoan mitoribosome. However, a superposition of the mitoribosome:LRPPRC-SLIRP-mRNA complex with *E. coli* expressome ^8^ shows not only that the nascent mRNA follows a comparable path in both systems, but also the mRNA delivering complexes bind in a similar location with respect to their ribosomes (Fig. 5b). To test whether protein-protein interactions can explain the conservation, we compared the interface with the *E. coli* expressome ^8^. Indeed, in the expressome, NusG binds to uS10 and restrains RNAP motions ^8^, and in our structure uS10m has a related interface between its α2 helix with mS31-mS39, which induces association of these two proteins (Fig. 5c, Extended Data Fig. 7).

Yet, most of the interactions rely on a mitochondria-specific N-terminal extension of uS10m where it shares a sheet with mS39 via the strand β1, and helices α1,16 and 18 are further involved in the binding (Extended Data Fig. 7). A similar conclusion can be reached from comparison with the *M. pneumoniae* expressome ^10^. Together, this analysis suggests that a specific protein-based mechanism must have evolved in the evolution of the metazoan mitoribosome for mRNA recognition and protection.

## Discussion

LRPPRC is an mRNA chaperone that regulates human mitochondrial transcription and translation and is involved in a neurodegenerative disorder. In this study, we report the cryo-EM structure of the LRPPRC-SLIRP in complex with the mitoribosome and characterize its function with respect to the mRNA delivery. We identified that LRPPRC, in complex with SLIRP, binds to mRNAs to hand-off transcripts to the mitoribosome for translation. The docking of LRPPRC is realized through the mitoribosomal proteins mS39 and the N-terminus of mS31, that together recognize eight of the LRPPRC helical repeats. The structural comparison with the unbound state uncovers that the N-terminus of mS31 adopts a stable conformation upon LRPPRC association.

Our structure also shows that SLIRP is directly involved in interactions with mRNA. Those interactions are supported by the comparison with other RNA-binding proteins that contain RNP domains, similar to SLIRP. SLIRP further stabilizes the architecture of LRPPRC, and both are required for mRNA binding. The mRNA is then channeled through a corridor formed with mS39 towards the decoding center.

Although *LRPPRC* knockout results in an overall decrease in the steady-state levels of the four OXPHOS complexes that contain mtDNA-encoded subunits, by implementing an RNAseq approach and metabolic labeling assays, we show that beyond its role in mRNA stabilization, LRPPRC has differential effects on the translational efficiency of mitochondrial transcripts.

Specifically, the synthesis of ND1, ND2, ATP6, COX1, COX2, and COX3 are particularly affected. Furthermore, our mitoribosome profiling data together with the structural analysis show that LRPPRC-SLIRP does not preexist on the mitoribosome as a structural element. Thus, the LRPPRC-SLIRP-dependent translation is not the sole regulatory pathway, and other mechanisms involving mRNA binding are likely to co-exist.

Since mS39 and mS31 are specific to Metazoa, as well as LRPPRC-SLIRP, the proposed mechanism in which some of the mitochondrial mRNAs are recruited for translation has developed in a co-evolutionary manner in Metazoa. However, also the presence of large RNA-binding moieties was also reported in association with mitoribosomes in other species ^56–60^.

Therefore, the principle of regulation by facilitation of molecular coupling might be a general feature, while unique molecular connectors involved in different species.

Overall, these findings define LRPPRC-SLIRP as a regulator of mitochondrial gene expression and explain how its components modulate the function of translation via mRNA binding. Given the challenge of studying mitochondrial translation due to the lack of an *in vitro* system, the native structures are crucial for explaining fundamental mechanisms. The identification of the components involved enhances our understanding of mitochondrial translation. Together, these studies provide the structural basis for translation regulation and activation in mitochondria.

## Methods

### Experimental model and culturing

HEK293S-derived cells (T501) were grown in Freestyle 293 Expression Medium containing 5% tetracycline-free fetal bovine serum (FBS) in vented shaking flasks at 37°C, 5% CO2 and 120 rpm (550 x g). Culture was scaled up sequentially, by inoculating at 1.5 x 10^6^ cells/mL and subsequently splitting at a cell density of 3.0 x 10^6^ cells/mL. Finally, a final volume of 2 L of cell culture at a cell density of 4.5 x 10^6^ cells/mL was used for mitochondria isolation, as previously described ^61^.

### Mitoribosome purification

HEK293S-derived cells were harvested from the 2 L culture when the cell density was 4.2 x 10^6^ cells/mL by centrifugation at 1,000 g for 7 min, 4 °C. The pellet was washed and resuspended in 200 mL Phosphate Buffered Saline (PBS). The washed cells were pelleted at 1,000 g for 10 min at 4 °C. The resulting pellet was resuspended in 120 mL of MIB buffer (50 mM HEPES-KOH, pH 7.5, 10 mM KCl, 1.5 mM MgCl2, 1 mM EDTA, 1 mM EGTA, 1 mM dithiothreitol, cOmplete EDTA-free protease inhibitor cocktail (Roche) and allowed to swell in the buffer for 15 min in the cold room by gentle stirring. About 45 mL of SM4 buffer (840 mM mannitol, 280 mM sucrose, 50 mM HEPES-KOH, pH 7.5, 10 mM KCl, 1.5 mM MgCl2, 1 mM EDTA, 1 mM EGTA, 1 mM DTT, 1X cOmplete EDTA-free protease inhibitor cocktail (Roche) was added to the cells in being stirred in MIB buffer and poured into a nitrogen cavitation device kept on ice. The cells were subjected to a pressure of 500 psi for 20 min before releasing the nitrogen from the chamber and collecting the lysate. The lysate was clarified by centrifugation at 800 x g and 4 °C, for 15 min, to separate the cell debris and nuclei. The supernatant was passed through a cheesecloth into a beaker kept on ice. The pellet was resuspended in half the previous volume of MIBSM buffer (3 volumes MIB buffer + 1 volume SM4 buffer) and homogenized with a Teflon/glass Dounce homogenizer. After clarification as described before, the resulting lysate was pooled with the previous batch of the lysate and subjected to centrifugation at 1,000 x g, 4 °C for 15 minutes to ensure complete removal of cell debris. The clarified and filtered supernatant was centrifuged at 10,000 x g and 4 °C for 15 min to pellet crude mitochondria. Crude mitochondria were resuspended in 10 mL MIBSM buffer and treated with 200 units of Rnase-free Dnase (Sigma-Aldrich) for 20 min in the cold room to remove contaminating genomic DNA. Crude mitochondria were again recovered by centrifugation at 10,000 g, 4 °C for 15 min and gently resuspended in 2 mL SEM buffer (250 mM sucrose, 20 mM HEPES-KOH, pH 7.5, 1 mM EDTA). Resuspended mitochondria were subjected to a sucrose density step-gradient (1.5 mL of 60% sucrose; 4 mL of 32% sucrose; 1.5 mL of 23% sucrose and 1.5 mL of 15% sucrose in 20 mM HEPES-KOH, pH 7.5, 1 mM EDTA) centrifugation in a Beckmann Coulter SW40 rotor at 28,000 rpm (139,000 x g) for 60 min. Mitochondria seen as a brown band at the interface of 32% and 60% sucrose layers were collected and snap-frozen using liquid nitrogen and transferred to −80 °C.

Frozen mitochondria were transferred on ice and allowed to thaw slowly. Lysis buffer (25 mM HEPES-KOH, pH 7.5, 50 mM KCl, 10 mM Mg(Oac)2, 2% polyethylene glycol octylphenyl ether, 2 mM DTT, 1 mg/mL EDTA-free protease inhibitors (Sigma-Aldrich) was added to mitochondria and the tube was inverted several times to ensure mixing. A small Teflon/glass Dounce homogenizer was used to homogenize mitochondria for efficient lysis. After incubation on ice for 5-10 min, the lysate was clarified by centrifugation at 30,000 x g for 20 min, 4 °C. The clarified lysate was carefully collected. Centrifugation was repeated to ensure complete clarification. A volume of 1 mL of the mitochondrial lysate was applied on top of 0.4 mL of 1 M sucrose (v/v ratio of 2.5:1) in thick-walled TLS55 tubes. Centrifugation was carried out at 231,500 x g for 45 min in a TLA120.2 rotor at 4 °C. The pellets thus obtained were washed and sequentially resuspended in a total volume of 100 µl resuspension buffer (20 mM HEPES-KOH, pH 7.5, 50 mM KCl, 10 mM Mg(Oac)2, 1% Triton X-100, 2 mM DTT). The sample was clarified twice by centrifugation at 18,000 g for 10 min at 4 °C. The sample was applied on to a linear 15-30% sucrose (20 mM HEPES-KOH, pH 7.5, 50 mM KCl, 10 mM Mg(Oac)2, 0.05% n-dodecyl-β-D-maltopyranoside, 2 mM DTT) gradient and centrifuged in a TLS55 rotor at 213,600 x g for 120 min at 4 °C. The gradient was fractionated into 50 μL volume aliquots. The absorption for each aliquot at 260 nm was measured and fractions corresponding to the monosome peak were collected. The pooled fractions were subjected to buffer exchange with the resuspension buffer.

### Cryo-EM data acquisition

3 μL of ∼120 nM mitoribosome was applied onto a glow-discharged (20 mA for 30 sec) holey-carbon grid (Quantifoil R2/2, copper, mesh 300) coated with continuous carbon (of ∼3 nm thickness) and incubated for 30 sec in a controlled environment of 100% humidity and 4 °C. The grids were blotted for 3 sec, followed by plunge-freezing in liquid ethane, using a Vitrobot MKIV (ThermoFischer). The data were collected on FEI Titan Krios (ThermoFischer) transmission electron microscope operated at 300 keV, using C2 aperture of 70 μm and a slit width of 20 eV on a GIF quantum energy filter (Gatan). A K2 Summit detector (Gatan) was used at a pixel size of 0.83 Å (magnification of 165,000X) with a dose of 29-32 electrons/Å^2^ fractionated over 20 frames.

### Cryo-EM data processing

The beam-induced motion correction and per-frame B-factor weighting were performed using RELION-3.0.2 ^62, 63^. Motion-corrected micrographs were used for contrast transfer function (CTF) estimation with gctf ^64^. Unusable micrographs were removed by manual inspection of the micrographs and their respective calculated CTF parameters. Particles were picked in RELION- 3.0.2, using reference-free followed by reference-aided particle picking procedures. Reference-free 2D classification was carried out to sort useful particles from falsely picked objects, which were then subjected to 3D classification. 3D classes corresponding to unaligned particles and LSU were discarded, and monosome particles were pooled and used for 3D auto-refinement yielding a map with an overall resolution of 2.9-3.4 Å for the five datasets. Resolution was estimated using a Fourier Shell Correlation cut-off of 0.143 between the two reconstructed half maps. Finally, the selected particles were subjected to per-particle defocus estimation, beam-tilt correction, and per-particle astigmatism correction followed by Bayesian polishing. Bayesian polished particles were subjected to a second round per-particle defocus correction. A total of 994,919 particles were pooled and separated into 86 optics groups in RELION-3.1 ^65^, based on acquisition areas and date of data collection. Beam-tilt, magnification anisotropy, and higher-order (trefoil and fourth-order) aberrations were corrected in RELION-3.1 ^65^. Particles with A-site and/or P-site occupied with tRNAs which showed comparatively higher occupancy for the unmodeled density potentially corresponding to the LRPPRC-SLIRP module, were pooled and re-extracted in a larger box size of 640 Å. The re-extracted particles were subjected to 3D autorefinement in RELION3.1 ^65^. This was followed by sequential signal subtraction to remove signal from the LSU, and all of the SSU except the region around mS39 and the unmodeled density, in that order. The subtracted data was subjected to masked 3D classification (T=200) to enrich for particles carrying the unmodeled density. Using a binary mask covering mS39 and all of the unmodeled density, we performed local-masked refinement on resulting 41,812 particles within an extracted sub-volume of 240 Å box size leading to 3.37 Å resolution map.

### Model building and refinement

At the mRNA channel entrance, a more accurate and complete model of mS39 could be built with 29 residues added to the structure. Improved local resolution enabled unambiguous assignment of residues to the density which allowed us to address errors in the previous model. A total of 28 α-helices could be modeled in their correct register and orientation. Further, a 28- residue long N-terminal loop of mS31 (residues 247–275) along mS39 and mito-specific N-terminal extension of uS9m (residues 53–70) approaching mRNA were modeled by fitting the loops in to the density maps.

For building LRPPRC-SLIRP module, the initial model of the full length LRPPRC was obtained from *AlphaFold2* Protein Structure Database (Uniprot ID P42704). Based on the analysis, three stable domains were identified that are connected by flexible linkers (673-983, 1035-1390). We then systematically assessed the domains against the map, and the N-terminal region (77-660) could be fitted into the density. The initial model was real space refined into the 3.37 Å resolution map of mS39-LRPPRC-SLIRP region obtained after partial signal subtraction using reference-restraints in *Coot* v0.9 ^66^. The N-terminal region covering residues 64-76 was identified in the density map and allowed us to model 34 helices of LRPPRC (residues 64-644). Helices α1-29 could be confidently modeled. An additional five helices, as predicted by *Alphafold2* ^34^, could be accommodated into the remaining density. After modelling LRPPRC into the map, there was an unaccounted density that fits SLIRP. The initial model of SLIRP was obtained from *Alphafold2* Protein Structure Database (Uniprot ID Q9GZT3). The unmodeled density agreed with the secondary structure of SLIRP. The model was real space refined into the density using reference-restraints as was done for LRPPRC in *Coot* v0.9 ^66^. Five additional RNA residues could be added to the 3’ terminal of mRNA to account for tubular density extending from it along the mRNA binding platform. The A/A P/P E/E state model was rigid body fitted into the corresponding 2.85 Å resolution consensus map. Modeled LRPPRC was merged with the rigid body fitted monosome model to obtain a single model of the mitoribosome bound to LRPPRC and SLIRP. The model was then refined against the composite map using PHENIX v1.18 ^67^ (Supplementary Information Table 1).

### Phylogenetic analysis

Phylogenetic distribution of proteins was determined by examining phylogeny databases ^55^, followed by sensitive homology detection to detect homologs outside of the bilateria. Orthologs were required to have identical domain compositions, and Dollo parsimony was used to infer the evolutionary origin of a protein from its phylogenetic distribution. When multiple homologs of a protein were detected in a species, a neighbor-joining phylogeny was constructed to assess monophyly of putative orthologs to the human protein. The short length of the SLIRP candidate protein from *T. adhaerens* (B3SAC0_TRIAD), that is part of the large RRM family, precludes obtaining a reliable phylogeny to confidently assess its orthology to human SLIRP is therefore tentative.

### TLSMD analysis

The TLSMD analysis ^35, 36^ was performed with full length LRPPRC model obtained from AlphaFold Database (AF-P42704-F1), and mitochondrial targeting sequence (residues 1-59) was removed. The model was divided into TLS segments (N), and single chain TLSMD is performed on all atoms using the isotropic analysis model. Instead of using atomic B-factors, the values for a per residue confidence score of AlphaFold called predicted local distance difference test (pLDDT) were used as reference to calculate the least squared residuals against the corresponding values calculated by TLSMD analysis. This is based on the assumption that local mobility of the model should be inversely correlated with the pLDDT score. AlphaFold pLDDT values and the corresponding calculated values were plotted for every iteration to monitor improvement in prediction and across the length of LRPPRC. The data in Extended Data Fig. 2 is presented for N=4, where segments 1 and 2 (residues 60-373 and 374-649) correspond to the modeled region, whereas segments 3 and 4 correspond to the remaining domains that could not be modeled.

### Helicase sequence analysis

To address the possibility that LRPPRC may serve as a helicase, we inspected the sequence of full-length LRPPRC (Uniprot ID P42704). First, we checked the sequence for matches with consensus motifs characteristic of helicases using regular expression search. The following motifs were searched, GFxxPxxIQ, AxxGxGKT, PTRELA, TPGR, DExD, SAT, FVxT, RgxD (DDX helicases); GxxGxGKT, TQPRRV, TDGML, DExH, SAT, FLTG, TNIAET, QrxGRAGR

(DHX helicases); AHTSAGKT, TSPIKALSNQ, MTTEIL (others). Next, we carried out multiple sequence analysis against representative member helicases of the DHX and DDX family to verify the results of the regular-expression sequence search and to find potentially valid weaker matches.

### Human cell lines and cell culture conditions

Human HEK293T embryonic kidney cells (CRL-3216, RRID: CVCL-0063) were obtained from ATCC. The HEK293T *LRPPRC* knock-out (KO) cell line was engineered in-house and previously reported ^27^. The LRPPRC-KO cell line was reconstituted with either the wild-type *LRPPRC* gene ^27^, or a variant causing Leigh syndrome, French-Canadian type (LSFC). The LSFC variant carries a single base change (nucleotide C1119T transition), predicting a missense A354V change at a conserved protein residue ^46^.

Cells were cultured in high-glucose Dulbecco’s modified Eagle’s medium (DMEM, Thermo Fisher Scientific, CAT 11965092), supplemented with 10% FBS (Thermo Fisher Scientific, CAT A3160402), 100 μg/mL of uridine (Sigma, CAT U3750), 3 mM sodium formate (Sigma CAT 247596), and 1 mM sodium pyruvate (Thermo Fisher Scientific, CAT 11360070) at 37 °C under 5% CO2. Cell lines were routinely tested for mycoplasma contamination.

To generate an *LRPPRC*-KO cell line reconstituted with the LSFC variant of the gene, a Myc-DDK tagged *LRPPRC* ORF plasmid was obtained from OriGene (CAT: RC216747). This ORF was then subcloned into a hygromycin resistance-containing pCMV6 entry vector (OriGene, CAT PS100024) and used to generate an *LRPPRC*-KO cell line reconstituted with a wild-type *LRPPRC* gene as reported ^27^. To generate the *LRPPRC*-LSFC variant carrying the C1119T mutation, we used the Q5® Site-Directed Mutagenesis Kit from NEB. ∼ 10 pg of template pCMV6-A-Myc-DDK-Hygro-*LRPPRC* vector were used, along with the primers LSFC-Q5-F 5’ GGAAGATGTAGTGTTGCAGATTTTAC and LSFC-Q5-R 5’ AATTTTTCAGTGACTAAAAGTAAAATG, designed to include the codon to be mutated. After exponential amplification and treatment with kinase and ligase, 2.5 µl of the reaction were transformed into competent *Escherichia coli* cells. Several transformants were selected, their plasmid DNA purified, and then sequenced to select the correct pCMV6-A-Myc-DDK-Hygro- *LRPPRC-LSFC* construct.

For transfection of the construct into *LRPPRC*-KO cells, we used 5 μl of EndoFectin mixed with 2 μg of vector DNA in OptiMEM-I media according to the manufacturer’s instructions. Media was supplemented with 200 μg/ml of hygromycin after 48 h, and drug selection was maintained for at least one month.

### Whole-Cell extracts and Mitochondria isolation

For SDS-PAGE electrophoresis, pelleted cells were solubilized in RIPA buffer (25 mM Tris-HCl pH 7.6, 150 mM NaCl, 1% NP-40, 1% sodium deoxycholate, and 0.1% SDS) with 1 mM PMSF (phenylmethylsulfonyl fluoride) and mammalian protease inhibitor cocktail (Sigma). Whole-cell extracts were cleared by 5 min centrifugation at 20,000 x *g* at 4 °C.

Mitochondria-enriched fractions were isolated from at least ten 80% confluent 15-cm plates as described previously ^68–70^. Briefly, the cells were resuspended in ice-cold T-K-Mg buffer (10 mM Tris–HCl, 10 mM KCl, 0.15 mM MgCl2, pH 7.0) and disrupted with 10 strokes in a homogenizer (Kimble/Kontes, Vineland, NJ). Using a 1 M sucrose solution, the homogenate was brought to a final concentration of 0.25 M sucrose. A post-nuclear supernatant was obtained by centrifugation of the samples twice for 5 min at 1,000 x *g*. Mitochondria were pelleted by centrifugation for 10 min at 10,000 x *g* and resuspended in a solution of 0.25 M sucrose, 20 mM Tris-HCl, 40 mM KCl, and 10 mM MgCl2, pH 7.4.

### Denaturing and native electrophoresis, followed by immunoblotting

Protein concentration was measured by the Lowry method ^71^. 40–80 μg of mitochondrial protein extract was separated by denaturing SDS–PAGE in the Laemmli buffer system ^72^. Then, proteins were transferred to nitrocellulose membranes and probed with specific primary antibodies against the following proteins: β-ACTIN (dilution 1:2,000; Proteintech; Rosemont, IL; 60008-1- Ig), ATP5A (1:1000; Abcam; Cambridge, MA; ab14748), CORE2 (1:1,000; Abcam; Cambridge,

MA; ab14745), COX1 (dilution 1:2,000; Abcam; Cambridge, MA; ab14705), LRPPRC (dilution 1:1,000; Proteintech; Rosemont, IL; 21175-1-AP), NDUFA9 (1:1000; Proteintech; Rosemont, IL; 20312-1-AP), SDHA (1:1,000; Proteintech; Rosemont, IL; 14865-1-AP) or SLIRP (1:1000; Abcam; Cambridge, MA; ab51523). Horseradish peroxidase-conjugated anti-mouse or anti-rabbit IgGs were used as secondary antibodies (dilution 1:10,000; Rockland; Limerick, PA). β- ACTIN was used as a loading control. Signals were detected by chemiluminescence incubation and exposure to X-ray film.

Blue-native polyacrylamide gel electrophoresis (BN-PAGE) analysis of mitochondrial OXPHOS complexes in native conditions was performed as described previously ^73, 74^. To extract mitochondrial proteins in native conditions, we pelleted and solubilized 400 μg mitochondria in 100 μl buffer containing 1.5 M aminocaproic acid and 50 mM Bis-Tris (pH 7.0) with 1% n-dodecyl b-D-maltoside (DDM). Solubilized samples were incubated on ice for 10 min in ice and pelleted at 20,000 x *g* for 30 min at 4 °C. The supernatant was supplemented with 10 µl of sample buffer 10X (750 mM aminocaproic acid, 50 mM Bis-Tris, 0.5 mM EDTA (ethylenediaminetetraacetic acid), 5% Serva Blue G-250). Native PAGE™ Novex® 3-12% Bis-Tris Protein Gels (Thermo Fisher) gels were loaded with 40 μg of mitochondrial proteins. After electrophoresis, the gel was stained with 0.25% Coomassie brilliant blue R250, or proteins were transferred to PVDF membranes using an eBlot L1 protein transfer system (GenScript, Piscataway, NJ) and used for immunoblotting.

### Pulse Labeling of Mitochondrial Translation Products

To determine mitochondrial protein synthesis, 6-well plates were pre-coated at 5 μg/cm^2^ with 50 μg/mL collagen in 20 mM acetic acid and seeded with WT or LRPPRC cell lines (two wells per sample per timepoint). 70% confluent cell cultures were incubated for 30 min in DMEM without methionine and then supplemented with 100 μl/ml emetine for 10 min to inhibit cytoplasmic protein synthesis as described ^68^. 100 μCi of [^35^S]-methionine was added and allowed to incorporate to newly synthesized mitochondrial proteins for increasing times from 15- to 60- minute pulses. Subsequently, whole-cell extracts were prepared by solubilization in RIPA buffer, and equal amounts of total cellular protein were loaded in each lane and separated by SDS-PAGE on a 17.5% polyacrylamide gel. Gels were transferred to a nitrocellulose membrane and exposed to a Kodak X-OMAT X-ray film. The membranes were then probed with a primary antibody against β-ACTIN as a loading control. Optical densities of the immunoreactive bands were measured using the Histogram function of the Adobe Photoshop software in digitalized images.

### Whole-cell transcriptomics

Cells were grown to 80% confluency in a 10 cm plate (two plates per sample) and were collected by trypsinization and washed once with PBS before resuspending in one mL of Trizol (ThermoFisher Scientific). RNA was extracted following the Trizol manufacturer’s specifications. The aqueous phase was transferred to a new tube, and an equal volume of 100% isopropanol and 3 μL of glycogen were added to precipitate the RNA. The sample was incubated at -80 °C overnight and centrifuged at 15,000 x*g* for 45 min at 4°C. RNA was resuspended in 50 μL of RNAse-free water and quantified by measuring absorbance at a wavelength of 260 nm. 2 μg of RNA was sent to Novogene (Sacramento, CA) for further processing. Novogen services included library preparation, RNA sequencing (RNAseq) on an Illumina HiSeq platform according to the Illumina Tru-Seq protocol, and bioinformatics analysis. The raw data was cleaned to remove low-quality reads and adapters using Novogen in-house Perl scripts in Cutadapt ^75^. The reads were mapped to the reference genome using the HISAT2 software ^76^. The transcripts were assembled and merged to obtain an mRNA expression profile with the StringTie algorithm ^77^, the RNA-seq data was then normalized to account for the total reads sequenced for each sample (the read depth), and differentially expressed mRNAs were identified by using the Ballgown suite ^78^ and the DESeq2 R package ^79^. GraphPad Prism version 9.0 software (GraphPad Software, San Diego, CA, USA) was used to prepare the volcano plots.

### Mitoribosome profiling

Mitoribosome profiling, matched RNA-seq, and data analysis were performed as described ^29^. Briefly, human and mouse cell lysates were prepared and mixed 95:5 human:mouse. For mitoribosome profiling, the combined lysates were subjected to RNaseI treatment and fractionated across a linear sucrose gradient. Sequencing libraries were prepared from the monosome fraction after phenol/chloroform extraction. For RNA-seq, RNA was extracted from the undigested combined lysate, fragmented by alkaline hydrolysis, and sequencing libraries prepared. Reads were cleaned of adapters, filtered of rRNA fragments, and PCR duplicates were removed. Read counts were summed across features (coding sequences) using Rsubread feature Counts ^80^, then normalized by feature length and mouse spike-in read counts. TE was calculated by dividing spike-in normalized mitoRPF reads per kilobase by spike-in normalized RNA-seq reads per kilobase. Values are expressed as log2-fold change in the *LRPPRC*-KO cells compared to the *LRPPRC* rescue cells. Mitoribosome profiling and RNA-seq data for *LRPPRC*-KO and *LRPPRC*-reconstituted cell lines are deposited in GEO under the accession number GSE173283. Mitoribosome profiling and RNA-seq data for the LSFC-reconstituted is deposited in GEO under the accession number GSEXXXXXX.

The mitoRPF length distribution was determined from mitochondrial mRNA-aligned reads. First, soft-clipped bases were removed using jvarkit ^81^, then frequency for each length was output using Samtools stats ^82^.

## Data availability statement

The atomic coordinates were deposited in the RCSB Protein Data Bank, and EM maps have been deposited in the Electron Microscopy Data bank under accession numbers 8ANY and EMD- 15544.

The atomic coordinates that were used in this study: <u>6ZTJ</u> (*E.coli* 70S-RNAP expressome complex in NusG); <u>6ZTN</u> (*E.coli* 70S-RNAP expressome complex in NusG); <u>1RKJ</u> (human Nucleolin); <u>5WWE</u> (human hnRNPA2/B1); <u>1CVJ</u> (Poly-adenylate binding protein, PABP)

## Acknowledgments

We thank S. Aibara and J. Andrell for help with data collection. The research was funded by the Swedish Foundation for Strategic Research (FFL15:0325), Ragnar Söderberg Foundation (M44/16), European Research Council (ERC-2018-StG-805230), Knut and Alice Wallenberg Foundation (2018.0080), NIH R01 (R01-GM123002) to SC, NIH R35 (R35-GM118141) to AB. V.S. was supported by the Horizon 2020 - Marie Sklodowska-Curie Innovative Training Network (721757), Y.I. was supported by H2020-MSCA-IF-2017 (799399-Itohribo), and C.M was supported by the Eunice Kennedy Shriver National Institute Of Child Health & Human Development of the National Institutes of Health under Award Number F30HD107939. The SciLifeLab cryo-EM facility is funded by the Knut and Alice Wallenberg, Family Erling Persson, and Kempe foundations. The content is solely the responsibility of the authors and does not necessarily represent the official views of the National Institutes of Health.

## Author contributions

V.S. collected cryo-EM data, processed the data, and built the models. V.S., Y.I. and A.A. performed structural analysis. C.M., F.F. and A.B. performed mitochondrial translation, OXPHOS, and RNAseq analysis. I.S., M.C. and S.C. performed mitoribosome profiling and RNAseq analysis. V.S., M.H. and A.A. performed evolutionary analysis. A.A. wrote the manuscript. All authors contributed to data interpretation and manuscript writing.

---

**Extended Data Fig. 1.**
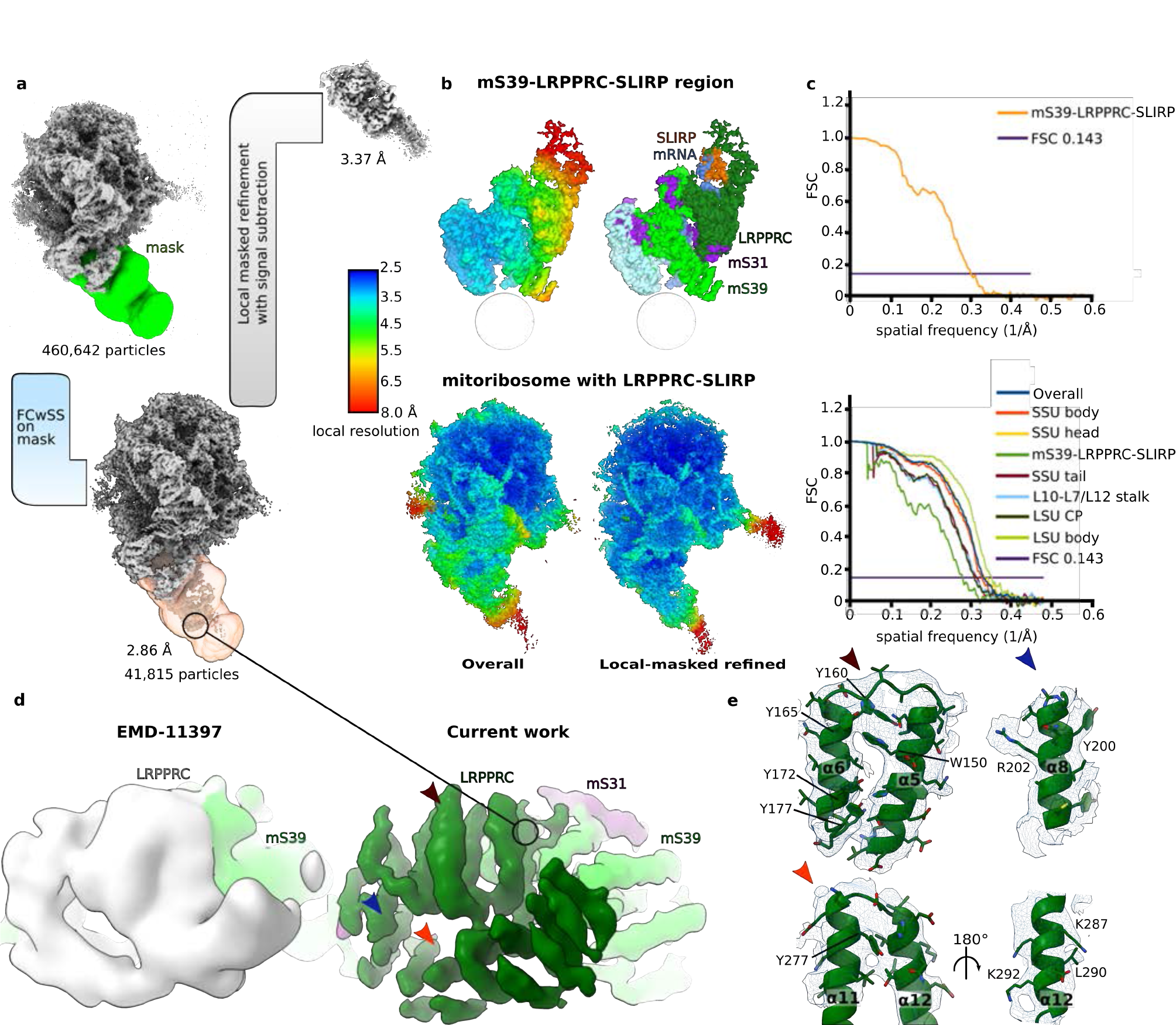
Cryo-EM data processing and map for mS39-LRPPRC-SLIRP region. **a.** Focused 3D-classification with signal subtraction using mask around mS39-LRPPRC-SLIRP region (transparent orange) of mitoribosome particles to identify LRPPRC-SLIRP containing monosome particles (2.86 Å overall resolution), followed by, masked refinement with signal subtraction on mS39-LRPPRC-SLIRP region to improve the local resolution. **b.** The mS39- LRPPRC-SLIRP map is shown colored by local resolution (top left) and by proteins assigned to the density (top right). The consensus map (bottom left) and the masked refined maps shown as a single composite map colored by local resolution (bottom right). **c.** Fourier shell correlation curves for the post-subtraction masked refined mS39-LRPPRC-SLIRP map (top) and individual masked refined maps. **d.** Map comparison for LRPPRC region between our work and EMD-11397. The map has been Gaussian filtered for better visibility. **e.** Density shown as mesh around helices α5-6, 8 and 11-12. Corresponding regions are indicated with arrows in panel (d).

**Extended Data Fig. 2.**
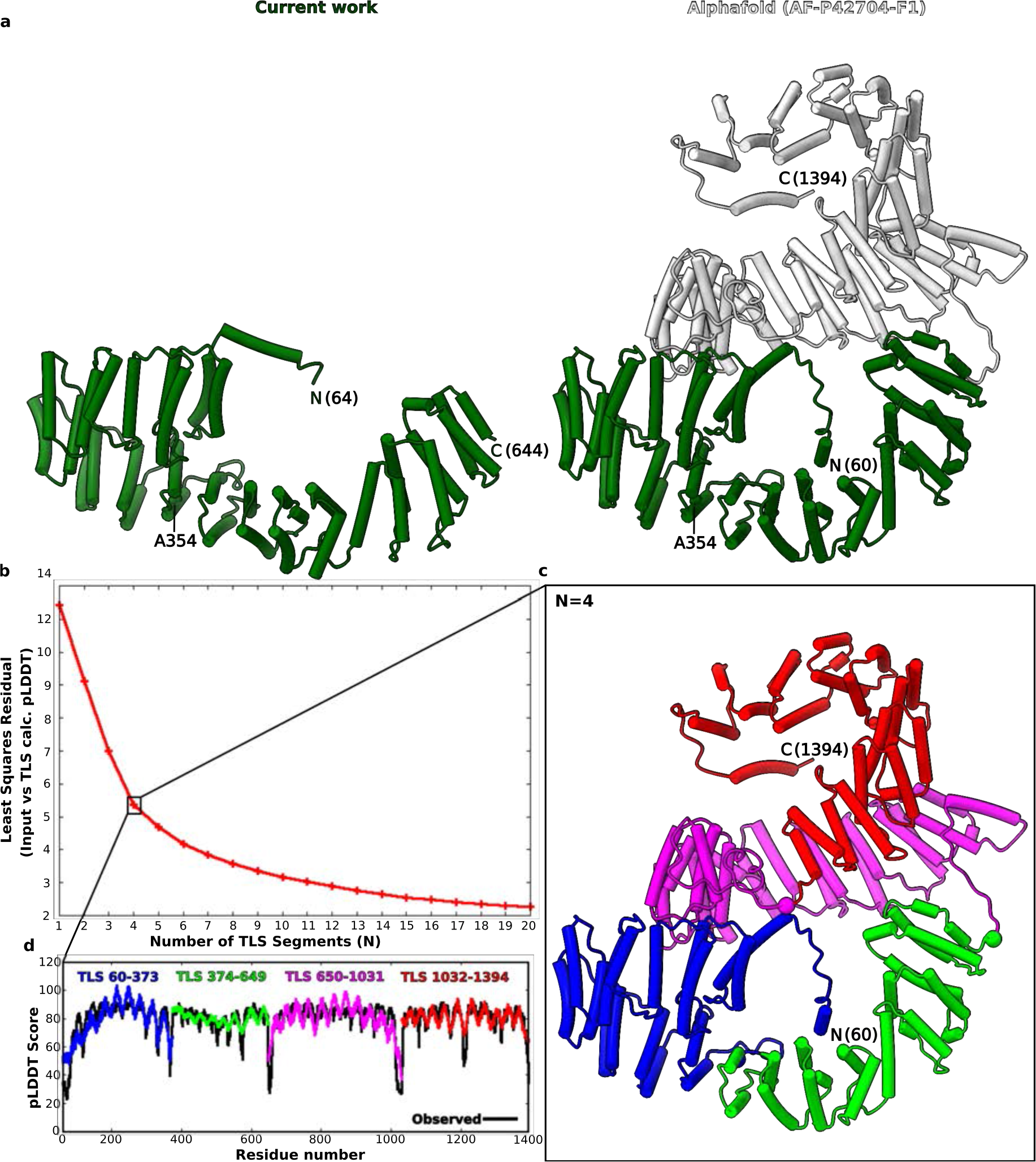
*AlphaFold* model and TLSMD analysis of LRPPRC. **a.** The modeled region of LRPPRC (residues 64-644) is compared with the AlphaFold model (AF- P42704-F1) of full length (right). The modeled region is green, the unmodeled is white. The position of LSFC variant (A354V) is indicated. **b.** TLSMD analysis of the AlphaFold model of LRPPRC up to 20 TLS segments (N). Graph plots least-square residuals assigned per-residue confidence score values (pLDDT) versus those calculated by TLS analysis. **c.** Model colored by TLS segments for N=4. Regions between the segments with high pLDDT values correspond to loop regions and are shown as spheres **d.** Comparison of AlphaFold assigned versus calculated pLDDT values at N=4.

**Extended Data Fig. 3.**
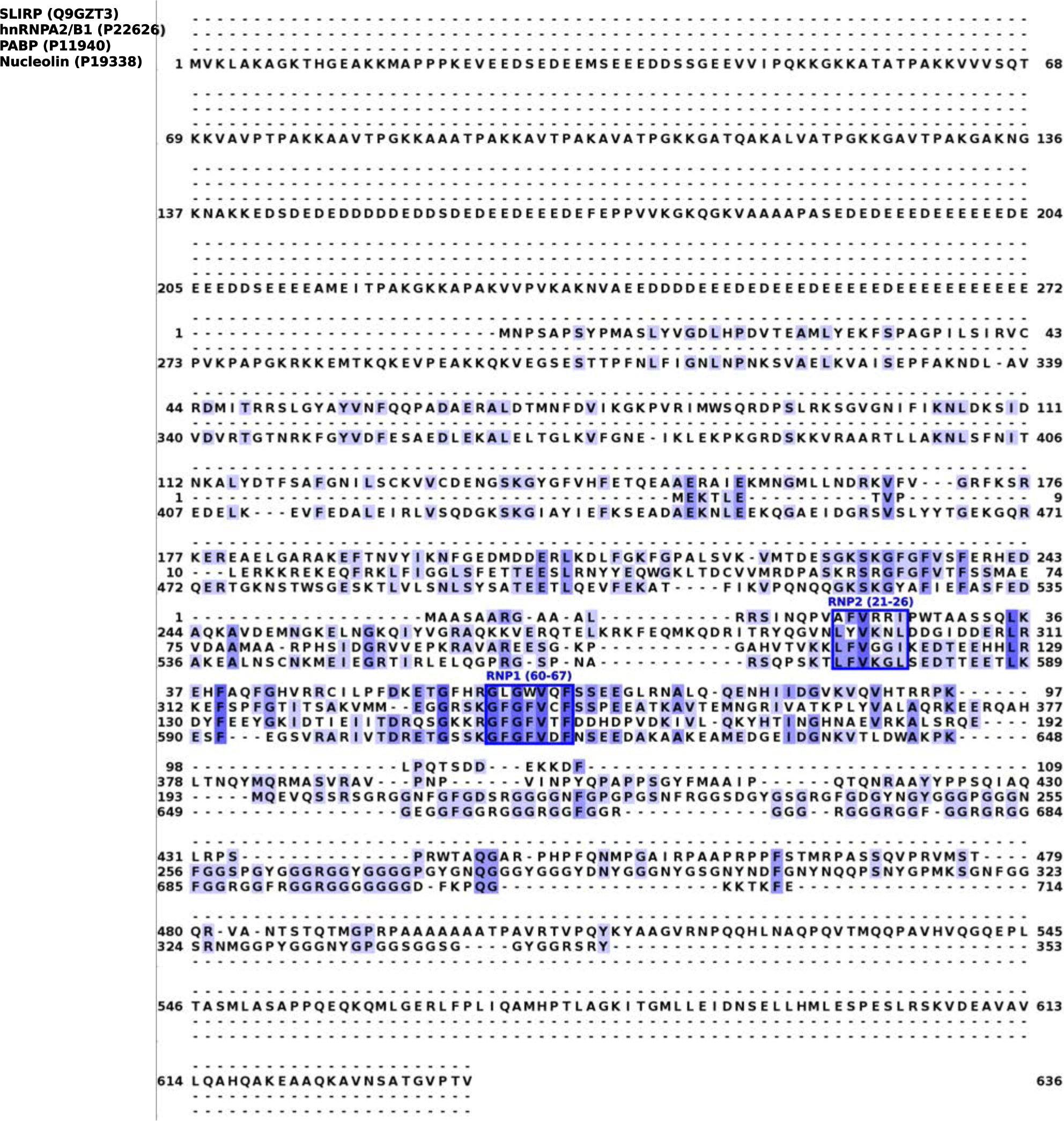
Multiple sequence alignment between SLIRP and representative RRM containing proteins. Alignment of SLIRP with representative RRM family proteins, heterogeneous nuclear ribnucleoproteins (hnRNPA2/B1), poly-A binding protein (PABP), and nucleolin shows conservation of submotifs RNP1 and RNP2 highlighted and indicated by corresponding residue numbers in SLIRP. Individual sequences are marked by residue numbers in the beginning and end and residues are colored by present identity.

**Extended Data Fig. 4.**
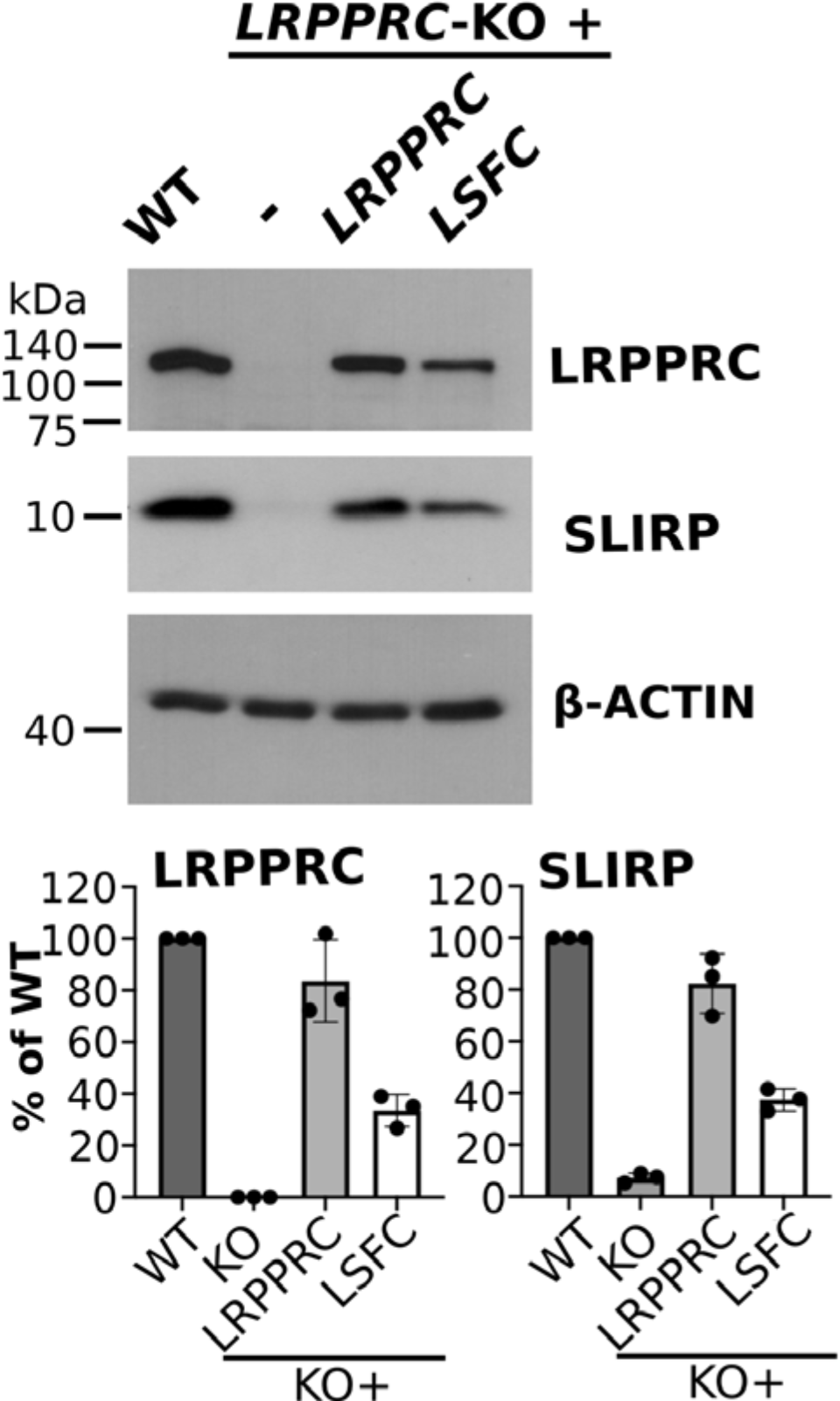
Reconstitution of the *LRPPRC*-KO with wild-type and LSCF variants of LRPPRC. Immunoblot analysis to estimate the steady-state levels of LRPPRC and SLIRP in the indicated cell lines. β-ACTIN was used as a loading control. The images were digitized, and the specific signals were quantified using the histogram function of Adobe Photoshop from three independent repetitions.

**Extended Data Fig. 5.**
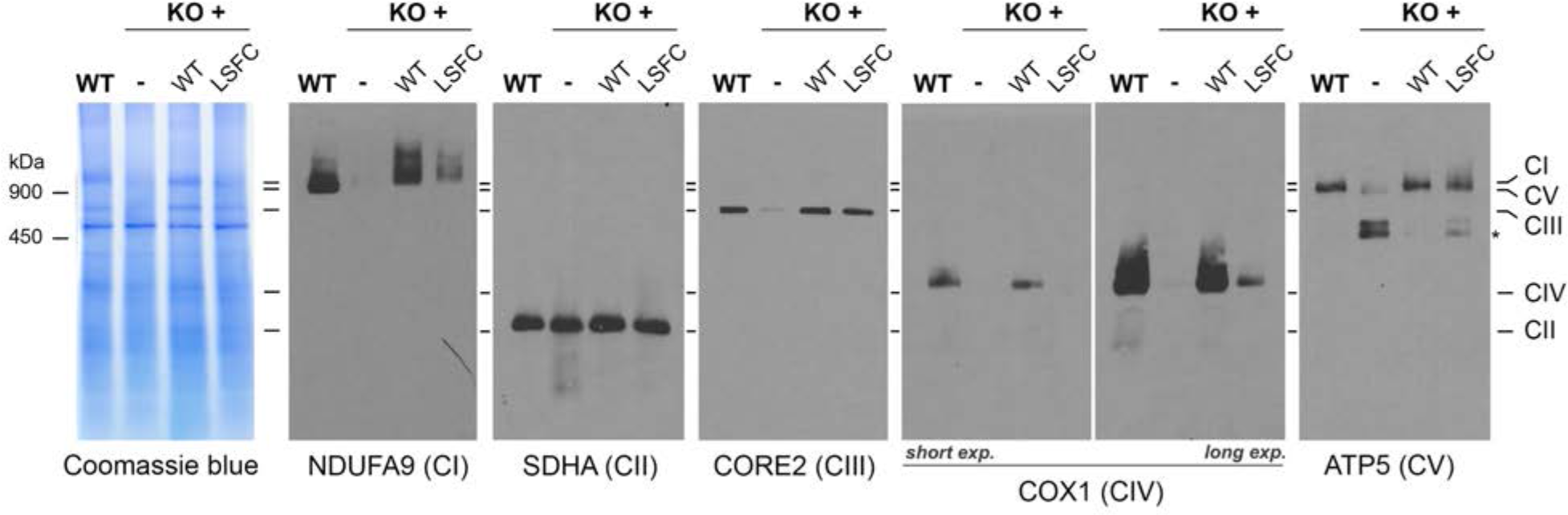
Mitochondrial protein synthesis is altered in *LRPPRC*-KO cells. Blue-native PAGE analyses in WT, *LRPPRC-*KO, and KO+WT cell lines. Intact respiratory complexes were extracted from purified mitochondria using 1% n-dodecyl β-D-maltoside. An asterisk indicates the ATPase (CV) F1 module that accumulates due to the low levels of the mitochondrion-encoded FO module subunits ATP6 and ATP8.

**Extended Data Fig. 6.**
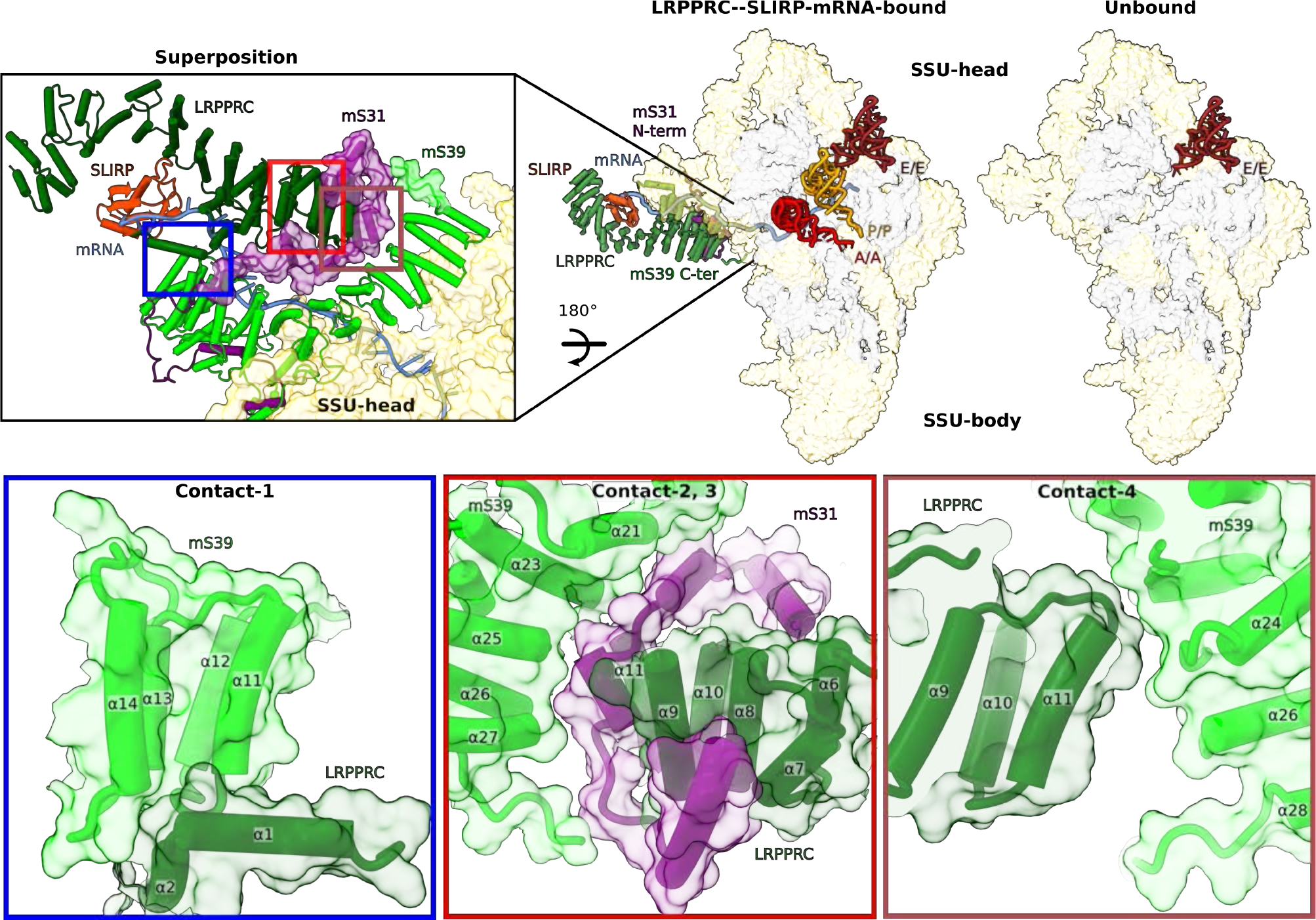
LRPPRC-SLIRP contacts with the SSU head. a. Comparison of SSU from mitoriboome:LRPPRC-SLIRP complex with SSU from E-site tRNA bound monosome. Zoom- in shows N-terminal region of mS31 and C-terminal loop of mS39 (in surface) stabilized by LRPPRC. **b.** Contact regions of LRPPRC with mS31 and mS39 shown in cartoon and surface representations.

**Extended Data Fig. 7.**
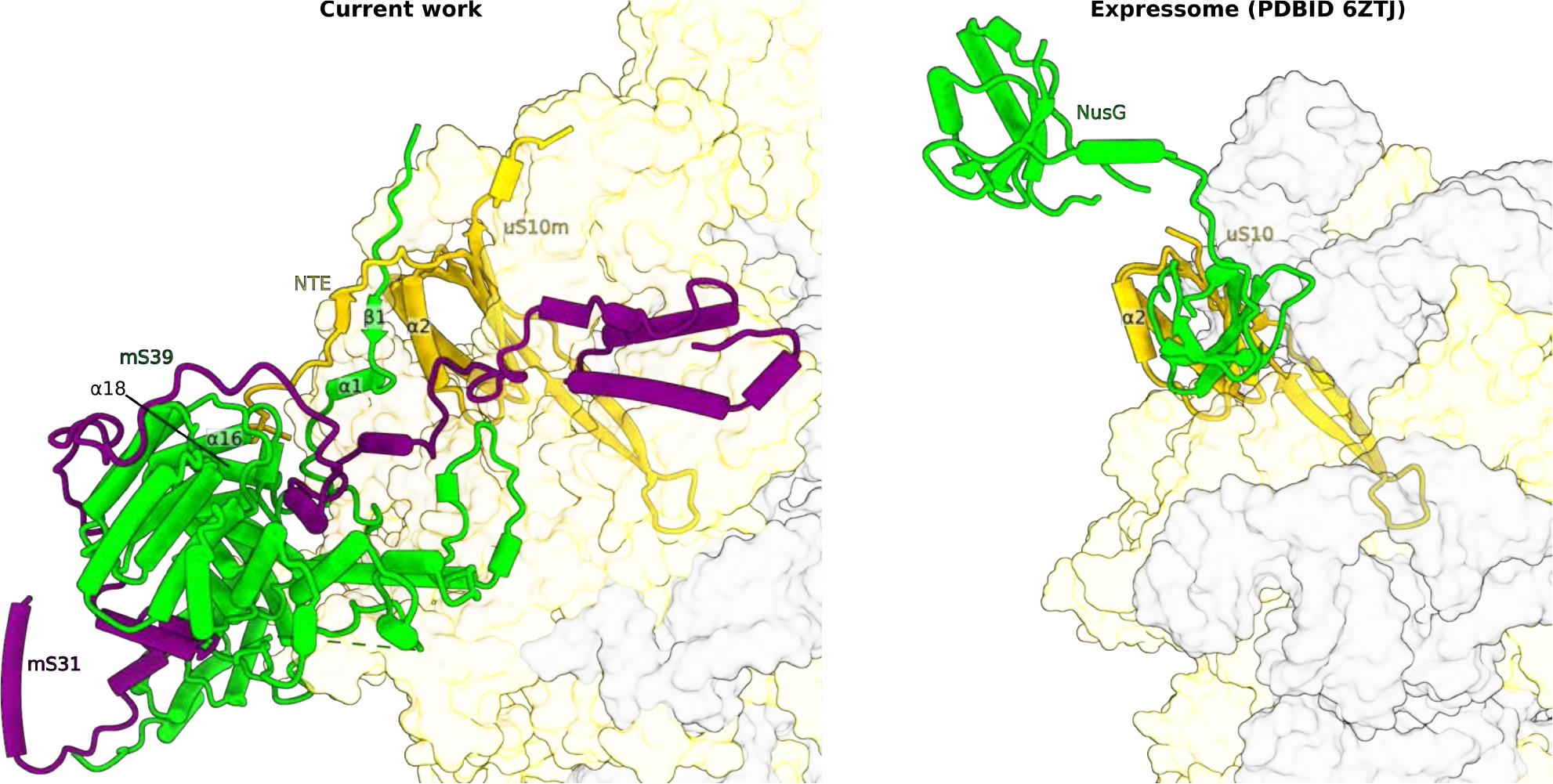
Close-up view of uS10m interactions with mS31-mS39. Interface between uS10m with mS31-mS39 that serve as the platform for LRPPRC-SLIRP is similar to that formed between uS10 and NusG that binds RNA polymerase in bacterial expressome

## SUPPLEMENTARY INFORMATION

**Supplementary Table 1.** Data collection and model statistics

|  | **Monosome with**<br>**LRPPRC:SLIRP**<br>**PDB: 8ANY**<br>**EMD-15544** |
| --- | --- |
| **Data collection and processing** |  |
| Electron microscope | Titan Krios |
| Camera | K2 Summit<br>(counting mode) |
| Magnification | 165,000 |
| Voltage (kV) | 300 |
| Electron exposure (e<sup>-</sup>/Å<sup>2</sup>) | 29-32 |
| No. of frames | 20 |
| Defocus range (μm) | -0.6 to -2.8 |
| Pixel size (Å) | 0.83 |
| Symmetry imposed | C<sub>1</sub> |
| Final particle number (no.) | 41,815 |
| Map resolution (Å) (Overall/<br>LSU-body/ CP/ L10-L12-stalk/ L1-stalk/ SSU-body/<br>SSU-head/ mS39/ SSU-tail/<br>mS39-LRPPRC-SLIRP) | 2.85/2.69/ 3.07/<br>3.07/ -/ 2.89/<br>2.84/-/ 3.02/3.37 |
| FSC threshold | 0.143 |
| Map resolution range (Å) | 2.5-8.0 |
| **Refinement** |  |
| Initial model used (PDB<br>code) | 6ZSG, 6RW4 |
| Model resolution (Å) | 2.7 |
| Model to map CC (CC<sub>volume</sub>) | 0.84 |
| FSC threshold | 0.5 |
| Map-sharpening *B* factor<br>(Å<sup>2</sup>) (Overall/ LSU-body/<br>CP/ L10-L12-stalk/ L1-stalk/<br>SSU-body/ SSU-head/ mS39/<br>Model composition) | -33/ -28/ -58/ -<br>49/-/ -39/ -39/-/-<br>55/ -60 |
| Non-hydrogen atoms | 356138 |
| Hydrogen atoms | 160529 |
| Protein chains | 90 |
| RNA chains | 7 |
| Protein residues (non-<br>modified/ *N*-acetylAla/ *N*-<br>acetylSer/ *N*-acetylThr *O*<sup>1</sup>-<br>methylisoAsp) | 15624/3/1/1/1 |
| RNA residues (non-<br>modified/ mG/ mU/ m<sup>1</sup>A/<br>m<sup>2</sup>G/ψ /m<sup>4</sup>C/ m<sup>5</sup>C/ m<sup>5</sup>U/<br>m<sup>6</sup>2A) | 2822/2/ 1/ 2/ 1/ 2/<br>1/ 1/ 1/ 2 |
| Ligands (ATP/ GDP/ NAD/<br>2Fe-2S/ spermine/<br>spermidine/ putrescine) | 1/ 1/ 1/ 3 /1 /4/ 1 |
| Ions (Zn<sup>2+</sup>/ K<sup>+</sup>/ Mg<sup>2+</sup>) | 3/ 49/ 206 |
| Waters | 6,926 |
| Mean atomic *B*-factor (Å<sup>2</sup>) |  |
| Protein | 38.28 |
| RNA | 36.42 |
| Ligand | 18.47 |
| Water | 15.21 |
| **Validation** |  |
| Ramachandran plot (%) |  |
| Outliers | 0.05 |
| Allowed | 1.85 |
| Favored | 98.11 |
| Clash score | 2.64 |
| RMSD |  |
| Bonds (Å) | 0.002 |
| Angles (°) | 0.432 |
| Rotamer outliers (%) | 0.00 |
| C<sub>β</sub> outliers (%) | 0.00 |
| CaBLAM outliers (%) | 0.90 |

**SI Video 1. Structure of LRPPRC-SLIRP module bound to monosome**

**Supplementary Video 1. Structure of LRPPRC-SLIRP module bound to monosome.**

The video shows the structure of LRPPRC-SLIRP determined in this work and how docking of mRNA on SSU is achieved by LRPPRC-SLIRP together with mito-specific proteins mS31 and mS39.

## References

1. Brown, A., Rathore, S., Kimanius, D., Aibara, S., Bai, X.-chen, Rorbach, J., Amunts, A., & Ramakrishnan, V. Structures of the human mitochondrial ribosome in native states of Assembly. Nature Structural & Molecular Biology, 24(10), 866–869 (2017).

2. Sissler, M., & Hashem, Y. Mitoribosome assembly comes into view. Nature Structural & Molecular Biology, 28(8), 631–633 (2021).

3. Itoh, Y. et al. Mechanism of mitoribosomal small subunit biogenesis and preinitiation. Nature 606, 603–608 (2022).

4. Lavdovskaia, E., Hillen, H. S., & Richter-Dennerlein, R. Hierarchical folding of the catalytic core during mitochondrial ribosome biogenesis. Trends in Cell Biology, 32(3), 182–185 (2022).

5. Harper, N.J., Burnside, C. & Klinge, S. Principles of mitoribosomal small subunit assembly in eukaryotes. Nature 614, 175–181 (2023).

6. Khawaja, A. et al. Distinct pre-initiation steps in human mitochondrial translation. Nat Commun 11, 2932, (2020).

7. Rackham, O., Filipovska, A. Organization and expression of the mammalian mitochondrial genome. Nat Rev Genet. 10, 606–623 (2022).

8. Webster, M. W. et al. Structural basis of transcription-translation coupling and collision in bacteria. Science 369, 1355–1359 (2020).

9. Wang, C. et al. Structural basis of transcription-translation coupling. Science 369, 1359– 1365 (2020).

10. O’Reilly, F. J. et al. In-cell architecture of an actively transcribing-translating expressome.Science 369, 554–557 (2020).

11. Itoh, Y. et al. Mechanism of membrane-tethered mitochondrial protein synthesis. Science 371, 846–849 (2021).

12. Ott, M., Amunts, A. & Brown, A. Organization and regulation of mitochondrial protein synthesis. Annual Review of Biochemistry 85, 77–101 (2016).

13. Christian, B. and Spremulli, L. Preferential selection of the 5′-terminal start codon on leaderless mRNAs by mammalian mitochondrial ribosomes J Biol Chem 285, 28379–28386 (2010).

14. Yi, SH. et al. Conformational rearrangements upon start codon recognition in human 48S translation initiation complex. Nucleic Acid Research 50, 5282–5298 (2022).

15. Lapointe, C.P. et al. eIF5B and eIF1A reorient initiator tRNA to allow ribosomal subunit joining. Nature, 607 (7917) 185–190 (2022).

16. Ruzzenente, B. et al. LRPPRC is necessary for polyadenylation and coordination of translation of mitochondrial mrnas. The EMBO Journal 31, 443–456 (2012).

17. Siira, S. J. et al. LRPPRC-mediated folding of the mitochondrial transcriptome. Nature Communications 8, (2017).

18. Chujo, T. et al. LRPPRC/SLIRP suppresses pnpase-mediated mrna decay and promotes polyadenylation in human mitochondria. Nucleic Acids Research 40, 8033–8047 (2012).

19. Sasarman, F., Brunel-Guitton, C., Antonicka, H., Wai, T. & Shoubridge, E. A. LRPPRC and SLIRP interact in a ribonucleoprotein complex that regulates posttranscriptional gene expression in mitochondria. Molecular Biology of the Cell 21, 1315–1323 (2010).

20. Mootha, V. K. et al. Identification of a gene causing human cytochrome *c* oxidase deficiency by Integrative Genomics. Proceedings of the National Academy of Sciences 100, 605–610 (2003).

21. Antonicka, H. et al. A high-density human mitochondrial proximity interaction network. Cell Metab 32, 479–497 (2020).

22. Sasarman, F., Brunel-Guitton, C., Antonicka, H., Wai, T. & Shoubridge, E.A. LRPPRC and SLIRP interact in a ribonucleoprotein complex that regulates posttranscriptional gene expression in mitochondria. Mol. Biol. Cell 21, 1315–1323. (2010).

23. Lagouge, M. et al. SLIRP regulates the rate of mitochondrial protein synthesis and protects LRPPRC from degradation. PLOS Genetics 11, (2015).

24. Baughman, J. M. et al. A computational screen for regulators of oxidative phosphorylation implicates SLIRP in mitochondrial RNA homeostasis. PLoS Genetics 5, (2009).

25. Guo, L. et al. Pathogenic SLIRP variants as a novel cause of autosomal recessive mitochondrial encephalomyopathy with complex I and IV deficiency. European Journal of Human Genetics 29, 1789–1795 (2021).

26. Sasarman, F. et al. Tissue-specific responses to the LRPPRC founder mutation in French Canadian leigh syndrome. Human Molecular Genetics 24, 480–491 (2014).

27. Xu, F., Addis, J. B. L., Cameron, J. M. & Robinson, B. H. LRPPRC mutation suppresses cytochrome oxidase activity by altering mitochondrial RNA transcript stability in a mouse model. Biochemical Journal 441, 275–283 (2011).

28. Spåhr, H. et al. SLIRP stabilizes LRPPRC via an RRM–PPR protein interface. Nucleic Acids Research 44, 6868–6882 (2016).

29. Soto, I. et al. Balanced mitochondrial and cytosolic translatomes underlie the biogenesis of human respiratory complexes. Genome Biol. 23, 170 (2022).

30. Aibara, S., Singh, V., Modelska, A. & Amunts, A. Structural basis of mitochondrial translation. eLife 9, (2020).

31. Amunts, A., Brown, A., Toots, J., Scheres, S. H. & Ramakrishnan, V. The structure of the human mitochondrial ribosome. Science 348, 95–98 (2015).

32. Greber, B. J. et al. The complete structure of the 55S mammalian mitochondrial ribosome. Science 348, 303–308 (2015).

33. Itoh, Y., et al. Structure of the mitoribosomal small subunit with streptomycin reveals Fe-S clusters and physiological molecules eLife 11:e77460 (2022).

34. Jumper J, et al. Highly accurate protein structure prediction with Alphafold. Nature 596, 583–589 (2021).

35. Painter, Jay, and Ethan A. Merritt. “Optimal description of a protein structure in terms of multiple groups undergoing TLS motion.” Acta Crystallographica Section D: Biological Crystallography 62, 439–450 (2006).

36. Painter, Jay, and Ethan A. Merritt. “TLSMD web server for the generation of multi-group TLS models.” Journal of applied crystallography 39, 109–111 (2006).

37. 37. Coquille, S. & Thore, S. Leigh syndrome-inducing mutations affect LRPPRC / SLIRP complex formation. *bioRxiv* https://doi.org/10.1101/2020.04.16.044412 (2020).

38. Burd, C. G., & Dreyfuss, G. Conserved structures and diversity of functions of RNA- binding proteins. Science, 265, 615–621 (1994).

39. Hatchell, E. C., Colley, S. M., Beveridge, D. J., Epis, M. R., Stuart, L. M., Giles, K. M., … & Leedman, P. J. SLIRP, a small SRA binding protein, is a nuclear receptor corepressor. Molecular cell, 22, 657–668 (2006).

40. Johansson, C., Finger, L. D., Trantirek, L., Mueller, T. D., Kim, S., Laird-Offringa, I. A., & Feigon, J. Solution structure of the complex formed by the two N-terminal RNA-binding domains of nucleolin and a pre-rRNA target. Journal of molecular biology, 337, 799–816 (2004).

41. Williams, P., Li, L., Dong, X., & Wang, Y. Identification of SLIRP as a G quadruplex- binding protein. Journal of the American Chemical Society, 139, 12426–12429 (2017).

42. Deo, R. C., Bonanno, J. B., Sonenberg, N., & Burley, S. K. Recognition of polyadenylate RNA by the poly (A)-binding protein. Cell, 98, 835–845 (1999).

43. Wu, B., Su, S., Patil, D. P., Liu, H., Gan, J., Jaffrey, S. R., & Ma, J. Molecular basis for the specific and multivariant recognitions of RNA substrates by human hnRNP A2/B1. Nature communications, 9, 1–12 (2018).

44. Takyar, S., Hickerson, R. P., & Noller, H. F. mRNA helicase activity of the ribosome. Cell, 120, 49–58 (2005).

45. Umate, P., Tuteja, N., & Tuteja, R. Genome-wide comprehensive analysis of human helicases. Communicative & Integrative Biology, 4, 118–137 (2011).

46. Kummer E, Leibundgut M, Rackham O, Lee RG, Boehringer D, Filipovska A & Ban N. Unique features of mammalian mitochondrial translation initiation revealed by cryo-EM Nature 560, 263–267 (2018).

47. Xu, F., Morin, C., Mitchell, G., Ackerley, C. & Robinson, B.H. The role of the *LRPPRC* (leucine-rich pentatricopeptide repeat cassette) gene in cytochrome oxidase assembly: mutation causes lowered levels of COX (cytochrome c oxidase) I and *COX III* mRNA. Biochem J. 382, 331–336. (2004).

48. Ruzzenente, B. et al. LRPPRC is necessary for polyadenylation and coordination of translation of mitochondrial mRNAs. EMBO J. 31, 443–456 (2012).

49. Couvillion, M.T., Soto, I.C., Shipkovenska, G. & Churchman, L.S. Synchronized mitochondrial and cytosolic translation programs. Nature 533, 499–503 (2016).

50. Ingolia, N.T., Ghaemmaghami, S., Newman, J.R. & Weissman, J.S. Genome-wide analysis in vivo of translation with nucleotide resolution using ribosome profiling. Science 324, 218–223 (2009).

51. Petrov, A. S. et al. Structural patching fosters divergence of mitochondrial ribosomes. Molecular Biology and Evolution 36, 207–219 (2018).

52. Huerta-Cepas, J. et al. Eggnog 5.0: A hierarchical, functionally and phylogenetically annotated orthology resource based on 5090 organisms and 2502 viruses. Nucleic Acids Research 47, (2018).

53. Sterky, F. H., Ruzzenente, B., Gustafsson, C. M., Samuelsson, T. & Larsson, N.-G. LRPPRC is a mitochondrial matrix protein that is conserved in metazoans. Biochemical and Biophysical Research Communications 398, 759–764 (2010).

54. Eddy, S. R. Accelerated profile HMM searches. PLoS Computational Biology 7, (2011).

55. Erwin, D. H. Early metazoan life: divergence, environment and ecology. Philosophical Transactions of the Royal Society B: Biological Sciences, 370, 1–7 (2015).

56. Amunts, A. et al. Structure of the yeast mitochondrial large ribosomal subunit. Science 343, 1485–1489 (2014).

57. Tobiasson, V. & Amunts, A. Ciliate mitoribosome illuminates evolutionary steps of mitochondrial translation. eLife 9, (2020).

58. Waltz, F., Soufari, H., Bochler, A., Giegé, P. & Hashem, Y. Cryo-EM structure of the RNA- rich plant mitochondrial ribosome. Nature Plants 6, 377–383 (2020).

59. Waltz, F. et al. How to build a ribosome from RNA fragments in Chlamydomonas mitochondria. Nature Communications 12, (2021).

60. Tobiasson, V., Berzina, I. & Amunts, A. Structure of a mitochondrial ribosome with fragmented rRNA in complex with membrane-targeting elements. Nature Communications 13, 6132 (2022).

## Additional References

61. Aibara, S., Andréll, J., Singh, V. & Amunts, A. Rapid isolation of the mitoribosome from Hek Cells. Journal of Visualized Experiments (2018). doi:10.3791/57877.

62. Zivanov, J. et al. New tools for automated high-resolution cryo-EM structure determination in relion-3. eLife 7, (2018).

63. Zivanov, J., Nakane, T. & Scheres, S. H. A bayesian approach to beam-induced motion correction in Cryo-EM single-particle analysis. IUCrJ 6, 5–17 (2019).

64. Zhang, K. GCTF: Real-time CTF determination and correction. Journal of Structural Biology 193, 1–12 (2016).

65. Zivanov, J., Nakane, T. & Scheres, S. H. Estimation of high-order aberrations and anisotropic magnification from Cryo-EM data sets in *relion*-3.1. IUCrJ 7, 253–267 (2020).

66. Emsley, P. & Cowtan, K. *Coot*: Model-building tools for Molecular Graphics. Acta Crystallographica Section D Biological Crystallography 60, 2126–2132 (2004).

67. Afonine, P. V. et al. Real-space refinement in*phenix*for cryo-EM and Crystallography. Acta Crystallographica Section D Structural Biology 74, 531–544 (2018).

68. Bourens, M., Boulet, A., Leary, S.C. & Barrientos, A. Human COX20 cooperates with SCO1 and SCO2 to mature COX2 and promote the assembly of cytochrome *c* oxidase. Hum. Mol. Genet. 23, 2901–2913 (2014).

69. Fernandez-Vizarra, E. et al. Isolation of mitochondria for biogenetical studies: An update. Mitochondrion 10, 253–262 (2010).

70. Moreno-Lastres, D. et al. Mitochondrial Complex I Plays an Essential Role in Human Respirasome Assembly. Cell Metab. 15, 324–335 (2012).

71. Lowry, O.H., Rosebrough, N.J., Farr, A.L. & Randall, R.J. Protein measurement with the Folin phenol reagent. J. Biol. Chem. 193, 265–275 (1951).

72. Laemmli, U.K. Cleavage of structural proteins during the assembly of the head of bacteriophage T4. Nature 227, 680–685 (1970).

73. Diaz, F., Barrientos, A. & Fontanesi, F. Evaluation of the mitochondrial respiratory chain and oxidative phosphorylation system using blue native gel electrophoresis. *Curr. Protoc. Hum. Genet.* Chapter, Unit19.14. (2009).

74. Timón-Gómez, A. et al. Protocol for the analysis of yeast and human mitochondrial respiratory chain complexes and supercomplexes by Blue Native-PAGE. STAR Protocols 1, 100119 (2020).

75. Martin, M. Cutadapt removes adapter sequences from high-throughput sequencing reads. EMBnet J. 17, 10–12 (2021).

76. Kim, D., Paggi, J.M., Park, C., Bennett, C. & Salzberg, S.L. Graph-based genome alignment and genotyping with HISAT2 and HISAT-genotype. Nat. Biotechnol. 37, 907–915 (2019).

77. Pertea, M. et al. StringTie enables improved reconstruction of a transcriptome from RNA- seq reads. Nat. Biotechnol. 33, 290–295 (2015).

78. Frazee, A.C. et al. Ballgown bridges the gap between transcriptome assembly and expression analysis. Nat. Biotechnol. 33, 243–246 (2015).

79. Love, M.I., Huber, W. & Anders, S. Moderated estimation of fold change and dispersion for RNA-seq data with DESeq2. Genome Biol. 15, 550 (2014).

80. Liao, Y., Smyth, G.K. & Shi, W. The R package Rsubread is easier, faster, cheaper and better for alignment and quantification of RNA sequencing reads. Nucleic Acids Res. 47, e47 (2019).

81. Lindenbaum, P. & Redon, R. bioalcidae, samjs and vcffilterjs: object-oriented formatters and filters for bioinformatics files. Bioinformatics 34, 1224–1225 (2018).

82. Danecek, P. et al. Twelve years of SAMtools and BCFtools. Gigascience 10 (2021).

